# Dual Membrane-spanning Anti-Sigma Factors Regulate Vesiculation in Gut Bacteroidota

**DOI:** 10.1101/2023.07.13.548920

**Authors:** Evan J. Pardue, Mariana G. Sartorio, Biswanath Jana, Nichollas E. Scott, Wandy Beatty, Juan C. Ortiz-Marquez, Tim Van Opijnen, Fong-Fu Hsu, Robert Potter, Mario F. Feldman

## Abstract

Bacteroidota are abundant members of the human gut microbiota that shape the enteric landscape by modulating host immunity and degrading dietary- and host-derived glycans. These processes are at least partially mediated by <u>O</u>uter <u>M</u>embrane <u>V</u>esicles (OMVs). In this work, we developed a high-throughput screen to identify genes required for OMV biogenesis and its regulation in *Bacteroides thetaiotaomicron* (*Bt*). Our screening led us to the identification of a novel family of <u>D</u>ual <u>M</u>embrane-spanning <u>A</u>nti-sigma factors (Dma), which regulate OMV biogenesis in *Bt*. We employed molecular and multiomic analyses to demonstrate that deletion of Dma1, the founding member of the Dma family, results in hypervesiculation by modulating the expression of NigD1, which belongs to a family of uncharacterized proteins found throughout Bacteroidota. Dma1 has an unprecedented domain organization: it contains a C-terminal β-barrel embedded in the OM; its N-terminal domain interacts with its cognate sigma factor in the cytoplasm, and both domains are tethered via an intrinsically disordered region that traverses the periplasm. Phylogenetic analyses reveal that the Dma family is a unique feature of Bacteroidota. This study provides the first mechanistic insights into the regulation of OMV biogenesis in human gut bacteria.

## Introduction

Vesiculation is a process by which cells utilize membranous compartments to traffic cellular contents (Deatherage and Cookson 2012; Sukhvinder et al. 2019). In eukaryotes, vesiculation, in the form of the trans-Golgi network, exosomes, and other extracellular vesicles, has been extensively studied (von Blume and Hausser 2019; Kalluri et al. 2020; Tricarico et al. 2017). However, much less is known about vesiculation in bacteria (Avila-Calderon et al. 2021; Kulp and Kuehn 2010; Sartorio et al. 2021). <u>O</u>uter <u>M</u>embrane <u>V</u>esicles (OMVs) are small, spherical structures generated by blebbing of the outer membrane (OM) in Gram-negative bacteria and therefore, they are comprised of OM components, like lipopolysaccharides (LPS), phospholipids, OM proteins, and periplasmic contents (Avila-Calderon et al. 2021; Kulp and Kuehn 2010; Sartorio et al. 2021). OMVs have been studied in many Gram-negative bacteria and are reported to mediate key bacterial processes, like pathogenesis, quorum sensing, immunomodulation, nutrient uptake, and envelope stress response (Bomberger et al. 2009; Rakoff-Nahoum et al. 2014; Durant et al. 2020; Hickey et al. 2015; Shen et al. 2012; Cooke et al. 2020; McBroom and Kuehn 2007). Vesicles can derive from active processes or by phenomena involving membrane destabilization. For example, recent reports have shown that explosive cell lysis is the primary source of vesicles in *Pseudomonas aeruginosa* (Turnbull et al. 2016), and it is likely that this process occurs in other closely related species (Toyofuku et al. 2023). In contrast, a growing body of literature has demonstrated that OMV biogenesis in *Bacteroides spp*. is a highly regulated process. Nonetheless, the mechanism of OMV biogenesis in Bacteroidota remains poorly understood.

*Bacteroides spp*. are gut commensals that comprise ∼40% of the bacterial species in the human gastrointestinal tract (Eckburg et al. 2005; Faith et al. 2013; Wexler and Goodman 2017). *Bacteroides* shape the intestinal environment by producing immunogenic compounds and degrading dietary- and host-derived glycans (Wexler et al. 2007; Zafar and Saier 2021). Studies have shown that OMVs produced by *Bacteroides spp*. facilitate many of their functions in the gut (Rakoff-Nahoum et al. 2014; Sartorio et al. 2023; Durant et al. 2020; Hickey et al. 2015; Shen et al. 2012). Transmission electron microscopy (TEM) revealed that *Bacteroides thetaiotaomicron* (*Bt*) and *Bacteroides fragilis* (*Bf)* produce large quantities of OMVs of uniform size (Elhenawy et al. 2014; Valguarnera et al. 2016). Mass spectrometry (MS) analyses revealed that OMV protein cargo is primarily composed of lipoproteins that function as glycosidases or proteases, and this cargo is tailored according to the available glycan landscape (Valguarnera et al. 2016; Sartorio et al. 2023). OMV-enriched lipoproteins contain a negatively charged motif (S-D/E_3_), called the <u>L</u>ipoprotein <u>E</u>xport <u>S</u>equence (LES), that is absent from OM-retained lipoproteins (Lauber et al. 2016; Valguarnera et al. 2018; Sartorio et al. 2023). These features are unprecedented and distinguish *Bacteroides* OMVs from the vesicles isolated by other Gram-negative bacteria. By exploiting these characteristics, we previously labeled OM- and OMV-specific proteins with fluorescent markers to visualize OMV biogenesis in *Bt*. Fluorescence microscopy was employed to observe OMVs actively blebbing from the OM of *Bt* in the absence of cell lysis (Sartorio et al. 2023). Altogether, these findings demonstrate that OMV biogenesis in *Bacteroides spp*. is the result of an active, regulated process, and not the result of cell lysis, as has been suggested for other bacteria. In this work, we developed a high-throughput screen to identify components of the machinery involved in OMV biogenesis and its regulation in *Bt*. We identified and characterized a new family of structurally novel <u>D</u>ual <u>M</u>embrane-spanning <u>A</u>nti-sigma factors (Dma) and investigated their role in modulating OMV biogenesis in *Bt*.

## Materials and Methods

### Bacterial strains and Growth conditions

Strains, oligonucleotides, and plasmids are described in **Table S1** in the supplemental material. *Escherichia coli* was grown aerobically at 37°C in Luria-Bertani (LB) medium. *Bacteroides* strains were grown in an anaerobic chamber (Coy Laboratories) at 37°C containing an atmosphere of 10% H_2_, 5% CO_2_, 85% N_2_. *Bacteroides* were cultured in Brain heart infusion (BHI) medium (Fisher Scientific) supplemented with 5 µg/ml Hemin and 1 µg/ml vitamin K3. When applicable, antibiotics were used as follows: 100 µg/ml ampicillin, 200 µg/ml gentamicin, 25 µg/ml erythromycin, and 10 μg/ml tetracycline.

### Genetic Manipulation of *Bt*

Deletion mutants were constructed using the pSIE1 vector described in Bencivenga-Barry et. al., 2020. Briefly, ∼750 base pair regions flanking genes of interest were cloned into pSIE1. Vectors were conjugated into *Bt* and positive conjugants were identified by selection on BHI plates containing gentamicin and erythromycin. Counterselection was performed on BHI plates in the presence or absence of 125 ng/mL anhydrotetracycline (aTc). Mutants were identified by PCR prior to genomic sequencing. Complemented strains were made using vectors from Whitaker et. al., 2017.

### OMV isolation

OMVs were purified by ultracentrifugation from cell-free culture supernatants. Briefly, 50mL of *Bt* culture grown to late stationary phase were centrifuged twice at 6,500 rpm at 4°C for 10 minutes. Supernatants were filtered using a 0.22-µm-pore membrane (Millipore) to remove residual cells. The filtrate was subjected to ultracentrifugation at 200,000xg for 2 h (Optima L-100 XP ultracentrifuge; Beckman Coulter). Supernatants were discarded, and pellets were resuspended in PBS standardized to OD_600_. When performing MS analysis, purified OMV preparations were lyophilized.

### Subcellular fractionation

<u>T</u>otal <u>M</u>embrane (TM) preparations were isolated by cell lysis and ultracentrifugation. Briefly, late stationary phase cultures were harvested by centrifugation at 6,500 rpm at 4°C for 10 minutes. The pellets were gently resuspended in a mixture of <u>P</u>hosphate <u>B</u>uffered <u>S</u>aline (PBS) containing complete EDTA-free protease inhibitor mixture (Roche Applied Science) followed by cell disruption. Centrifugation at 8,500 rpm at 4 °C for 8 minutes was performed to remove unbroken cells. Total membranes were collected by ultracentrifugation at 200,000 xg for 1 h at 4 °C. Supernatants were discarded, and pellets were resuspended in PBS standardized to OD_600_. TM fractions were lyophilized for MS analysis.

### SDS-PAGE and Western blot analyses

Total membrane and vesicle fractions were analyzed by standard 10% Tris-glycine SDS-PAGE. Samples were normalized by OD_600,_ and equivalent volumes were loaded onto an SDS-PAGE gel. Coomassie Blue staining was employed to analyze protein profiles. When applicable, samples were transferred onto a nitrocellulose membrane for Western blot analysis. Membranes were blocked using Tris-buffered saline (TBS)-based Odyssey blocking solution (LI-COR). Primary antibodies used in this study were rabbit polyclonal anti-His (ThermoFisher) and mouse monoclonal anti-FLAG (Sigma). Secondary antibodies used were IRDye anti-rabbit 780 (LI-COR). Imaging was performed using an Odyssey CLx scanner (LI-COR).

### Lipopolysaccharide (LPS) Silver stain

Abundance of LPS was measured according to Tsai and Frasch 1982. Briefly, samples we ran on 20% SDS-PAGE, then gels were fixed overnight in 200 mL of 40% ethanol in 5% acetic acid. Next, gels were oxidized for 5min in 100mL of 0.7% fresh periodic acid in 40% ethanol and 5% acetic acid. Upon completion, gels underwent three washes (15 min each) in milliQ H_2_O. The gels were then stained for 10min in the dark with 28 mL 0.1M NaOH, 2mL NH_4_OH, 5mL 20% AgNO_3_, and 115 milliQ H_2_O. Gels underwent three additional washes prior to developing in 200 mL H_2_O with 10 mg Citric acid and 100 μL Formaldehyde.

### OMV Reporter screen

Transposon mutagenesis was performed by adapting the protocol of Veeranagouda et. al., 2013 on *Bt* constitutively expressing BACOVA_04502 (Inulinase; INL) fused to Nanoluciferase (NLuc) at the C-terminus. Briefly, INL-NLuc was cloned in the pWW3867 vector (Whitaker et. al., 2017) and expressed in *Bt*. Next, cells were conjugated with pSAM-Bt_Tet **(Table S1)** to create the transposon mutant library. Selection was performed on BHI agar plates containing 25 µg/ml erythromycin and 10 μg/ml tetracycline. Upon plating, individual colonies were isolated and transferred to 200 μl of BHI media in clear, round-bottom 96-well plates (Corning) and incubated for 24 h. After incubation, samples were sub-cultured and diluted (1:20) in the same volume of BHI media and transferred to a second 96-well plate for a 20 h incubation. Next, we measured the OD_600_ of cultures in each plate with a BioTek microplate reader. Plates then underwent centrifugation at 4000rpm to pellet cells. Supernatants were collected and transferred to 0.22 μm hydrophilic low protein binding 96-well filter plates (Millipore) and centrifuged at 4000 rpm to remove residual cells. Quantification of OMV production in 96-well plate format was done by performing Nano-Glo assays using the Nano-Glo Live Cell Assay System (Promega). Briefly, 100 μl of filtered supernatants were transferred to white bottom 96-well plates (Corning) and 20μl of Nano-Glo Live Cell Reagent was added to each well. Plates were shaken for 20 s in a BioTek microplate reader prior to quantifying luminescent output. Luminescence was normalized to OD_600_. Transposon mutants displaying a >1.5-fold increase relative to the wild-type were considered hypervesiculating strains, while those displaying <0.5-fold decrease were deemed hypovesiculating strains. Transposon insertions from candidates of interest were identified by genomic DNA extraction and sequencing.

### RNA Sequencing Sample Collection, Library Preparation, and Analysis

WT and ΔDma1 were grown overnight in BHI media before being diluted to the equivalent of OD 0.1 in 10 mL and grown anaerobically for 4 h at 37°C. Four individual overnight and 10-mL culture biological replicates were prepared. Cultures were normalized, and an amount of culture equivalent to an OD_600_ of 4.0 was pelleted for 90 s at 8,000 rpm. Pellets were resuspended on ice in 1 mL TRIzol (Invitrogen) with 10 μL of 5 mg/mL glycogen. Samples were flash frozen and stored at −80°C until extraction. Prior to extraction, samples were thawed on ice, then pelleted, and supernatants were treated with chloroform. RNA was extracted from the aqueous phase using the RNeasy minikit (Qiagen, Inc.), and RNA quality was checked by agarose gel electrophoresis and *A*_260_/*A*_280_ measurements. RNA was stored at −80°C with SUPERase-IN RNase inhibitor (Life Technologies) until library preparation.

RNA sequencing prep (RNA-Seq) was performed as previously described (Zhu et. al., 2020). Briefly, 400ng of total RNA from each sample was used for generating cDNA libraries following our RNAtag-Seq protocol. PCR amplified cDNA libraries were sequenced on an Illumina NextSeq500, obtaining a high-sequencing depth (over 7 million reads per sample). RNA-Seq data was analyzed using our *in-house* developed analysis pipeline *Aerobio*. Raw reads are demultiplexed by 5’ and 3’ indices, trimmed to 59 base pairs, and quality filtered (96% sequence quality>Q14). Filtered reads are mapped to the corresponding reference genomes using bowtie2 with the --very-sensitive option (-D 20 –R 3 –N 0 –L 20 –i S, 1, 0.50). Mapped reads are aggregated by feature Count and differential expression is calculated with DESeq2 (Love et. al., 2014; Liao et. al., 2014). In each pair-wise differential expression comparison, significant differential expression is filtered based on two criteria: |log2foldchange| > 1 and adjusted p-value (*padj*) <0.05. All differential expression (DE) comparisons are made between the WT and ΔDma1 mutants under the condition mentioned above. The reproducibility of the transcriptomic data was confirmed by an overall high Spearman correlation across biological replicates (R > 0.95). BioProject ID PRJNA994135.

### Sample preparation for Proteomic analysis

WT, ΔDas1, ΔDma1, and ΔDas1-Dma1 were grown overnight anaerobically in 2mL of BHI media prior to being diluted into 50 mL and grown for 20 h. Whole cells, total membranes, and vesicles were collected from each strain. Four individual biological replicates of each fraction were performed for each strain. Samples were lyophilized in preparation for MS analysis.

Lyophilized protein samples were solubilized in 4% SDS, 100mM Tris pH 8.5 by boiling them for 10 min at 95 °C. The protein concentrations were assessed using a bicinchoninic acid protein assay (Thermo Fisher Scientific) and 100µg of each biological replicate prepared for digestion using Micro S-traps (Protifi, USA) according to the manufacturer’s instructions. Briefly, samples were reduced with 10mM DTT for 10 mins at 95°C and then alkylated with 40mM IAA in the dark for 1 hour. Samples were acidified to 1.2% phosphoric acid and diluted with seven volumes of S-trap wash buffer (90% methanol, 100mM Tetraethylammonium bromide pH 7.1) before being loaded onto S-traps and washed 3 times with S-trap wash buffer. Samples were then incubated with Trypsin (1:100 protease:protein ratio, in 100mM Tetraethylammonium bromide pH 8.5) overnight at 37°C before being collected by centrifugation with washes of 100mM Tetraethylammonium bromide pH 8.5, followed by 0.2% formic acid followed by 0.2% formic acid / 50% acetonitrile. Samples were then dried down and further cleaned up using homemade C18 Stage 1,2 tips to ensure the removal of any particulate matter.

### Reverse phase Liquid chromatography–mass spectrometry

C18 purified digests were re-suspended in Buffer A* (2% acetonitrile, 0.01% trifluoroacetic acid) and separated on a Ultimate 3000 UPLC (Thermo Fisher Scientific) with a two-column chromatography set up composed of a PepMap100 C18 20 mm x 75 μm trap and a PepMap C18 500 mm x 75 μm analytical column (Thermo Fisher Scientific). Digests were loaded on to the trap column at 5 μL/min for 6 minutes with Buffer A (0.1% formic acid, 2% DMSO) then infused into a Orbitrap 480™ at 300 nl/minute via the analytical column. 95-minute runs were undertaken by altering the buffer composition from 2% Buffer B to 28% B over 70 minutes, then from 25% B to 40% B over 4 minutes, then from 40% B to 80% B over 3 minutes. The composition was held at 80% B for 2 minutes, and then dropped to 2% B over 1 minutes before being held at 2% B for another 10 minutes. The Orbitrap 480™ Mass Spectrometer was operated in a data-dependent mode automatically switching between the acquisition of a single Orbitrap MS scan (300-1600 m/z, maximal injection time of 25 ms, an AGC set to a maximum of 300% and a resolution of 120k) every 3 seconds and Orbitrap MS/MS HCD scans of precursors (using a stepped NCE of 28,32,40%, maximal injection time of 80 ms, an AGC set to a maximum of 300% and a resolution of 30k). Identification and LFQ analysis were accomplished using MaxQuant (v1.6.17.0)3. Data was searched against the *B. thetaiotaomicron* reference proteome (Uniprot: UP000001414) allowing for oxidation on Methionine. The LFQ and “Match Between Run” options were enabled to allow comparison between samples. Maxquant search results were processed using Perseus (version 1.6.0.7) 3 with missing values imputed based on the total observed protein intensities with a range of 0.3 σ and a downshift of 1.8 σ. Statistical analysis was undertaken in Perseus using two-tailed unpaired T-tests between groups.

### Data availability

The mass spectrometry proteomics data has been deposited in the Proteome Xchange accession: PXD043360.

### MS data analysis

Identification and LFQ analysis were accomplished using Max-Quant (v2.0.2.0) 8 using *Bacteroides thetaiotaomicron* VPI-5482 proteome (Uniprot: UP000001414) allowing for oxidation on Methionine. Prior to MaxQuant analysis dataset acquired on the Fusion Lumos were separated into individual FAIMS fractions using the FAIMS MzXML Generator9. The LFQ and “Match Between Run” options were enabled to allow comparison between samples. The resulting data files were processed using Perseus (v1.4.0.6)10 to filter proteins not observed in at least four biological replicates of a single group. ANOVA and Pearson correlation analyses were performed to compare groups. Predicted localization and topology analysis for proteins identified by MS were performed using UniProt11, PSORT12, SignalP13 and PULDB14.

### LC-MS analysis of lipids from TM and OMVs

WT, ΔDas1, and its complemented strain were grown overnight anaerobically in 5mL of BHI media prior to being diluted into 140 mL and grown for 20 h. TMs and OMVs were collected from each strain. Four individual biological replicates of each fraction were performed for each strain. Total lipids were extracted according to Bligh and Dyer chloroform:methanol method (Bligh and Dyer 1959). Briefly, 2 volumes of methanol, 1 volume of chloroform, and 0.8 volumes of Milli-Q water were added to 1 volume of PBS-resuspended sample in solvent-resistant glass tubes. Contents were mixed for 2 min by vortexing, and 1 volume of chloroform was added to the mixture. Samples were vortexed for another minute, then centrifugated for 5 min at 4,000 rpm. After centrifugation, the bottom phase (organic) was recovered using a glass Pasteur pipette and stored in solvent-sealed vials at −80°C until lipid analysis by LC-MS.

Untargeted LC/MS analyses were conducted on an Agilent 6550 A QTOF instrument with an Agilent 1290 high-performance liquid chromatograph (HPLC) with an autosampler, operated using Agilent MassHunter software (Santa Clara, CA, USA). Separation of the total lipid extracts was achieved using a Thermo Fisher (Waltham, MA, USA) BETASIL C_18_ column (100 × 2.1 mm, 5 μm) at a flow rate of 300 μL/min at room temperature. The mobile phase contained 5 mM ammonium formate (pH 5.0) both in solvent A, acetonitrile:water (60:40, vol/vol), and solvent B, isopropanol:acetonitrile (90:10, vol/vol). A gradient elution was applied in the following manner: 68% A, 0 to 1.5 min; 68 to 55% A, 1.5 to 4 min; 55 to 48% A, 4 to 5 min; 48 to 42% A, 5 to 8 min; 42 to 34% A, 8 to 11 min; 34 to 30% A, 11 to 14 min; 30 to 25% A, 14 to 18 min; 25 to 3% A, 18 to 23 min; 3 to 0% A, 25 to 30 min; 0% A, 30 to 35 min; 68% A, 35 to 40 min. Both the positive-ion and negative-ion electrospray ionization (ESI) MS scans were acquired in the mass range of 200 to 2,000 Da at a rate of 2 scans/min. High-resolution (*R* = 100,000 at *m*/*z* 400) mass spectrometric analyses of the lipid extracts were also conducted on a Thermo LTQ Orbitrap Velos. Lipids were loop injected into the ESI ion source using a built-in syringe pump which was set to continuously deliver a flow of 20 μL/min methanol with 0.5% NH_4_OH. The scanned mass spectra were recalibrated internally with a known mass, namely, 13:0/15:0 PE at *m*/*z* 634.4453. Linear ion trap (LIT) multistage MS (MS^n^) spectra were obtained for structural identification as described previously (Hsu et. al., 2007; Hsu and Turk 2008; Hsu 2016).

### Negative staining and analysis by transmission electron microscopy

For quantitative analyses at the ultrastructural level, 200 mesh formvar/carbon-coated copper grids (Ted Pella Inc., Redding CA) were coated with 50µg/ml poly-L-lysine (Sigma, St Louis, MO) for 10 min at 37^○^C. Excess fluid was removed, and grids were allowed to air dry. Poly-L-lysine coating allowed for even distribution of material across the grid. Bacterial OMVs were fixed with 1% glutaraldehyde (Ted Pella Inc.) and allowed to absorb onto freshly glow discharged poly-L-lysine-coated grids for 10 min. Grids were then washed in dH_2_O and stained with 1% aqueous uranyl acetate (Ted Pella Inc.) for 1 min. Excess liquid was gently wicked off and grids were allowed to air dry. Samples were viewed on a JEOL 1200EX transmission electron microscope (JEOL USA, Peabody, MA) equipped with an AMT 8-megapixel digital camera (Advanced Microscopy Techniques, Woburn, MA). Each OMV sample was processed in triplicate (3 grids). Ten random images were taken at a magnification of 25,000x from various areas of the grid with a total of 90 images for each sample, and the number of OMV in each image was quantified.

### Phylogenetic analysis of Dma family

Whole-genome fasta files were obtained from NCBI Assembly on 06/28/23. Genomes were annotated for open reading frames with prokka v1.14.6 using the command “prokka ${ID}_*.fna --outdir ${ID} --locustag ${ID} --mincontiglen 500 --prefix ${ID} --force --notrna --norrna” {24642063}. The core genome alignment of the 29 genomes was created using panaroo v1.2.10 on the .gff files from prokka with the command “panaroo -i ${indir}/*.gff -o ${outdir} --clean-mode moderate -a core -c .8 -f .5 --core_threshold -t 12 --search_radius 10 --refind_prop_match 100 -t ${SLURM_CPUS_PER_TASK}” {32698896}. Alignment of the 29 core genes identified by panaroo was performed using mafft v7 {23329690}. The core genome alignment from panaroo was constructed into a maximum likelihood phylogenetic tree using RAxML v8.2.12 with the command “raxmlHPC -s core_gene_alignment.aln -n EP_raxml -m GTRGAMMA -f a -T 4 -N 100 -p 12345 - x 54321” {24451623}. Identification of Dma1, Dma2, and Dma3 homologs outside of *B. thetaiotamicron* was performed with the NCBI tblastn webserver using the protein sequences for each gene as query and the whole-genome fasta files as the subject with 95% query cover threshold as a positive result for gene presence. AlphaFold structural predictions were used to confirm genes identified during the blast search. The newick file was viewed and analyzed with metadata on Dma1, Dma2, Dma3, related gene information in iTOL webserver{33885785}.

## Results

### Screening for genes involved in OMV biogenesis

Vesiculation has been studied in Gram-negative bacteria for ∼60 years; however, our understanding of OMV biogenesis is still in its infancy. To identify bacterial proteins involved in OMV biogenesis in *Bt*, we developed a high throughput assay to quantify OMV production *in vitro*. We constructed an OMV reporter consisting of *Bacteroides ovatus* inulinase (BACOVA_04502; INL), a protein previously shown to be enriched in OMVs (Elhenawy et al. 2014; Sartorio et al. 2023), fused to Nanoluciferase (NLuc) (England et al. 2016). Western blotting confirmed that the fusion protein was stable and almost exclusively present in OMVs **(Fig. 1A)**. Filtered supernatants from cells constitutively expressing INL-NLuc exhibited significantly higher luminescent output than the lysis control strain expressing cytoplasmic NLuc (cNLuc) **(Fig. 1B)**. By using this method, luminescence in culture supernatants can be used as an easily quantifiable proxy for OMV production. Next, we created a transposon library in the strain expressing the chromosomally encoded INL-NLuc. Since INL-NLuc is almost exclusively trafficked to OMVs, we expected that mutants displaying abnormal levels of NLuc activity in their supernatants corresponded to abnormal levels of OMV production. The strategy employed for the screening is described in **Figure 1C**. We screened ∼5300 colonies and detected several mutants displaying abnormally high or low levels of NLuc activity **(Fig. S1)**. Secondary screening with TEM, western blotting, and LPS and protein analysis were performed to validate potential candidates. The genomes of mutants displaying atypical levels of vesiculation were sequenced to identify the transposon insertion sites. These analyses identified several hyper- and hypo-vesiculating strains, summarized in **Table S2**.

**Figure 1:**
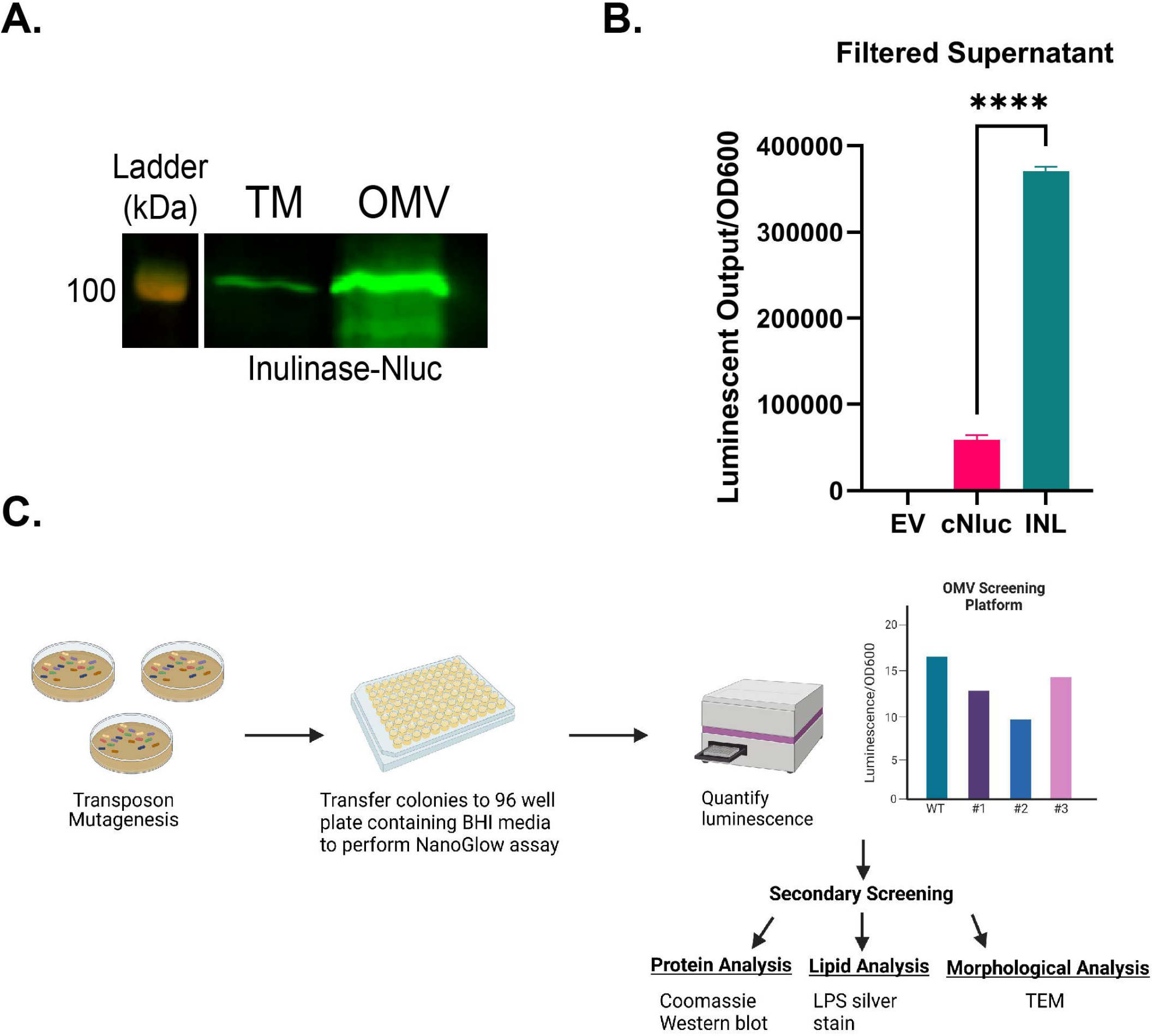
Identification of genes involved in OMV biogenesis. (A) Western blot against anti-polyHis showing TM and OMV fractions from BT WT constitutively expressing Inulinase fused to Nluc (INL-Nluc). This blot demonstrates that INL-Nluc is stably expressed and properly localizes to the OMV fraction. Samples were standardized by OD600 and ran on 10% SDS-PAGE. (B) Filtered supernatants from cells expressing INL-Nluc display higher luminescent output than those expressing Nluc in the cytoplasm. Filtered supernatants from overnight cultures (∼20hrs) were used for NanoGlow assays to quantify OMV production in vitro. Total luminescent output was normalized by OD_600_. (C) Schematic of OMV screening methodology (Created with BioRender.com).

### Mutation of Dma1 (BT_4721) leads to hypervesiculation

Genomic sequencing of potential candidates revealed four independent mutants containing transposon insertions in the gene BT_4721, which we renamed Dma1 **(Fig. 2A)**. Western blots showed that these mutants had increased abundance of INL-NLuc in the OMV fraction **(Fig. 2B)**, suggesting that Dma1 plays a role in OMV biogenesis. We generated a Dma1 deletion mutant (ΔDma1) and its corresponding complemented strain (ΔDma1_Comp_). Growth curves confirmed that the fitness of ΔDma1 is not attenuated in this context **(Fig. S2)**. Next, we isolated and quantified OMVs by TEM. This analysis confirmed that ΔDma1 produces significantly more OMVs than the WT **(Fig. 2C)**. *Bacteroides* OMVs contain lipopolysaccharide (LPS), membrane lipids (phospholipids, sphingolipids, and amino lipids), and protein cargo (Sartorio et. al., 2021). To validate the hypervesiculation phenotype biochemically, we measured the relative amounts of these components in total <u>m</u>embrane (inner and outer; TM) and OMV preparations from WT, ΔDma1, and ΔDma1_Comp_. We analyzed protein content by SDS-PAGE. Surprisingly, the electrophoretic profile of the OMV fraction from ΔDma1 exhibited a significant distortion in the low molecular weight region of the gel, which was reverted by complementation **(Fig. 2D)**. We hypothesized that the distortion was due to the presence of high amounts of LPS in the samples due to ΔDma1 hypervesiculation. Analysis of LPS from TM and OMV fractions via SDS-PAGE followed by silver stain confirmed that ΔDma1 secretes significantly more LPS than the wild-type and complemented strains **(Fig. 2E)**. The OMV fraction of ΔDma1 also contained more proteins than the wild-type and complemented strains **(Fig. 2F)**. We performed comparative proteomic analyses between the WT and ΔDma1 strains. Principal component analysis showed that WT and ΔDma1 TM and OMV contain similar protein content **(Fig. S3)**. More in-depth analysis revealed that ΔDma1 OMVs mostly carry lipoproteins containing the N-terminal LES motif, which constitute the proper OMV cargo in *Bt* (Sartorio et. al., 2023) **(Fig. S4A)**. Integral membrane proteins and lipoproteins lacking the LES domain were retained predominantly in the bacterial membranes. ΔDma1 OMVs did not contain significant amounts of inner membrane or cytoplasmic components, ruling out that the increase in the secreted amounts of LPS and proteins is due to cell lysis **(Fig. S4A)**. Finally, we performed comparative lipidomics on TM and OMV fractions from WT, ΔDma1, and ΔDma1_Comp_. While the lipid composition for both fractions remained relatively unperturbed, we detected a general increase in secreted phospho, sphingo-, and amino lipid content in the ΔDma1 OMVs **(Fig. 2G and Fig. S5)**. In summary, our microscopy, proteomic, lipidomic, and LPS analyses demonstrated that ΔDma1 produces significantly more bona fide OMVs than its parental strain.

**Figure 2:**
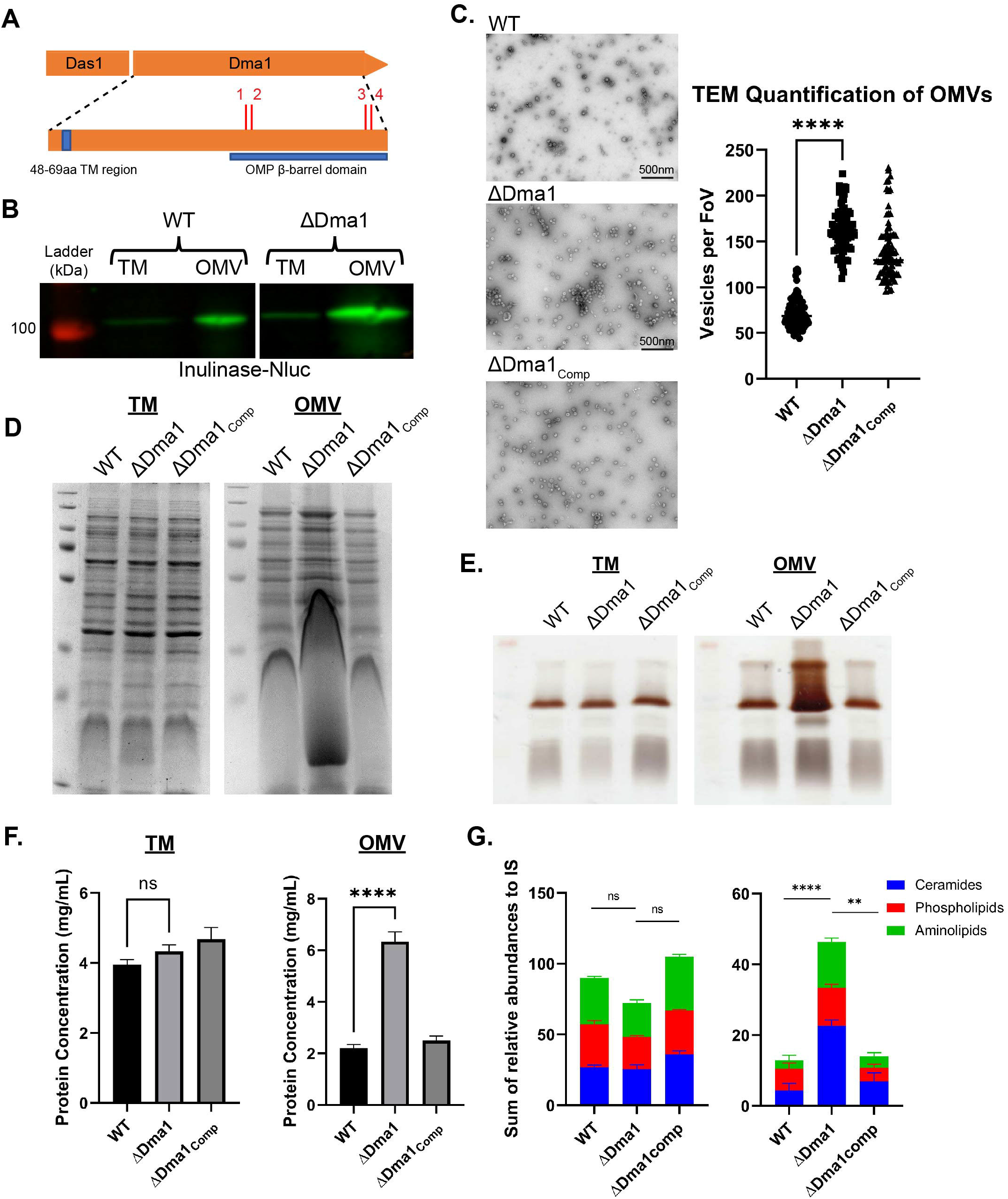
Deletion of Dma1 leads to hypervesiculation. (A) Genomic sequencing of screening candidates revealed four independent transposon mutants in Dma1. Red dashes denote transposon insertions (1: 674nt, 2: 677nt, 3: 1,139nt, 4: 1,143nt). (B) Western blot using anti-polyHis shows that Dma1 transposon mutants contain more INL-Nluc in the OMV fraction than the wild-type. Samples were normalized by OD_600_ and run on 10% SDS-PAGE. (C) Transmission electron microscopy (TEM) reveals that ΔDma1 produces significantly more OMVs than the wild-type. Left: Representative TEM images of OMVs from each strain. Right: Quantification of TEM images from each strain (FoV: Field of View). Three biological replicates of samples from each strain were fixed onto grids in triplicate (in materials and methods). Ten random images were taken from each grid (n=90 per strain) and OMVs were counted manually. (D) Coomassie Blue Stain (E) LPS Silver Stain demonstrate (F) Lowry Protein Assay each demonstrate that BTΔ4721 produces significantly more OMVs than the WT. (G) Total lipids were extracted from TMs and OMVs from WT, ΔDma1, and ΔDma1_Comp_ demonstrating that ΔDma1 secretes significantly more membrane lipids as a result of hypervesiculation. In all cases, samples were normalized to OD_600_.

### <u>E</u>xtra<u>c</u>ytoplasmic <u>F</u>unction (ECF) sigma factor, Das1 (BT_4720), is required for Dma1-mediated hypervesiculation

Dma1 is encoded in an operon with the ECF21 family sigma factor, BT_4720 (Das1, for <u>D</u>ma-<u>a</u>ssociated <u>s</u>igma factor 1. ECF-type sigma factors are a family of transcriptional regulators that modulate gene expression in response to extracellular signals. These are typically encoded adjacent to their cognate anti-sigma factor, which negatively regulates the activity of the sigma factor (Staron et al. 2009; Otero-Asman et al. 2019). ECF21 family sigma factors are found solely among Bacteroidota, but very little is known regarding their function (Staron et al. 2009). An ortholog of Dma1 in *Bf*, Reo, was shown to control the activity of its cognate sigma factor, ecfO, by sequestering it at the IM via its N-terminal region in the cytoplasm (Ndamukong et al. 2013). There is strong consensus between the amino acid sequences from Reo and Dma1 **(Fig. S6)**, which suggests that Dma1 possesses a cytoplasmic domain capable of directly regulating the activity of Das1. We deleted Das1 in the ΔDma1 background (ΔDas1-Dma1). A single deletion mutant lacking Das1 (ΔDas1) was included as control. By employing SDS-PAGE, we determined that removal of Das1 reverts the distortion in the electrophoretic pattern seen for ΔDma1 OMVs **(Fig. S7)**, indicating that the hypervesiculation observed in ΔDma1 requires Das1. Additional comparative proteomics confirmed that cargo selection was also conserved for each additional mutant **(Fig. S2, S3A-B)**. These findings demonstrate that Dma1 functions as the cognate anti-sigma factor for Das1, and indicates that the unregulated activity of Das1 is responsible for hypervesiculation in ΔDma1.

A previous study has shown that deletion of *Reo*, in *Bf*, increases fitness in response to oxidative stressors, while deletion of *ecfO* renders them more susceptible (Ndamukong et al. 2013). We assessed the fitness of WT, ΔDas1, ΔDma1, and their corresponding complemented strains in response to prolonged aerobic stress but observed no changes between the mutants and the wild-type **(Fig. S8)**. This suggests that Dma1 and *Reo* control different responses in *Bt* and *Bf*, respectively.

### NigD1, part of the Dma1-Das1 regulon, is necessary for hypervesiculation

To understand how Dma1 controls vesiculation, we performed RNA sequencing to compare the transcriptomes of wild-type and ∆Dma1 strains. We found that three genes, BT_1287, NigD1 (BT_4005), and NigD2 (BT_4719), were dramatically upregulated in the ∆Dma1 mutant **(Fig. 3A and Table S3)**. There were other genes differentially expressed **(Table S3)**, but their changes were much less pronounced. A comparative proteomic analysis of WT and ∆Dma1 identified the same three proteins as the most differentially expressed proteins between these strains **(Fig 3B and Table S4)**. No function has been assigned to any of these three proteins, however both, NigD1 and NigD2 are annotated as NigD-like proteins. NigD-like proteins are a family of uncharacterized lipoproteins proteins found solely in Bacteroidota. We determined that deletion of NigD1 (but not BT_1287 or NigD2) in the ∆Dma1 background restores wild-type levels of vesiculation, indicating that NigD1 is involved in the Dma1-controlled vesiculation process **(Fig. 3C)**. NigD1 is not encoded within an operon; however, it is encoded nearby genes required for LPS biosynthesis and regulation (LpxB and FtsH) and phospholipid synthesis (CdsA). Our transcriptomic and proteomic analyses failed to identify these, or any other known genes involved in the biosynthesis of LPS or phospholipid differentially regulated between WT and ΔDma1. However, our results suggest that NigD1 acts as a molecular switch for OMV biogenesis.

**Figure 3:**
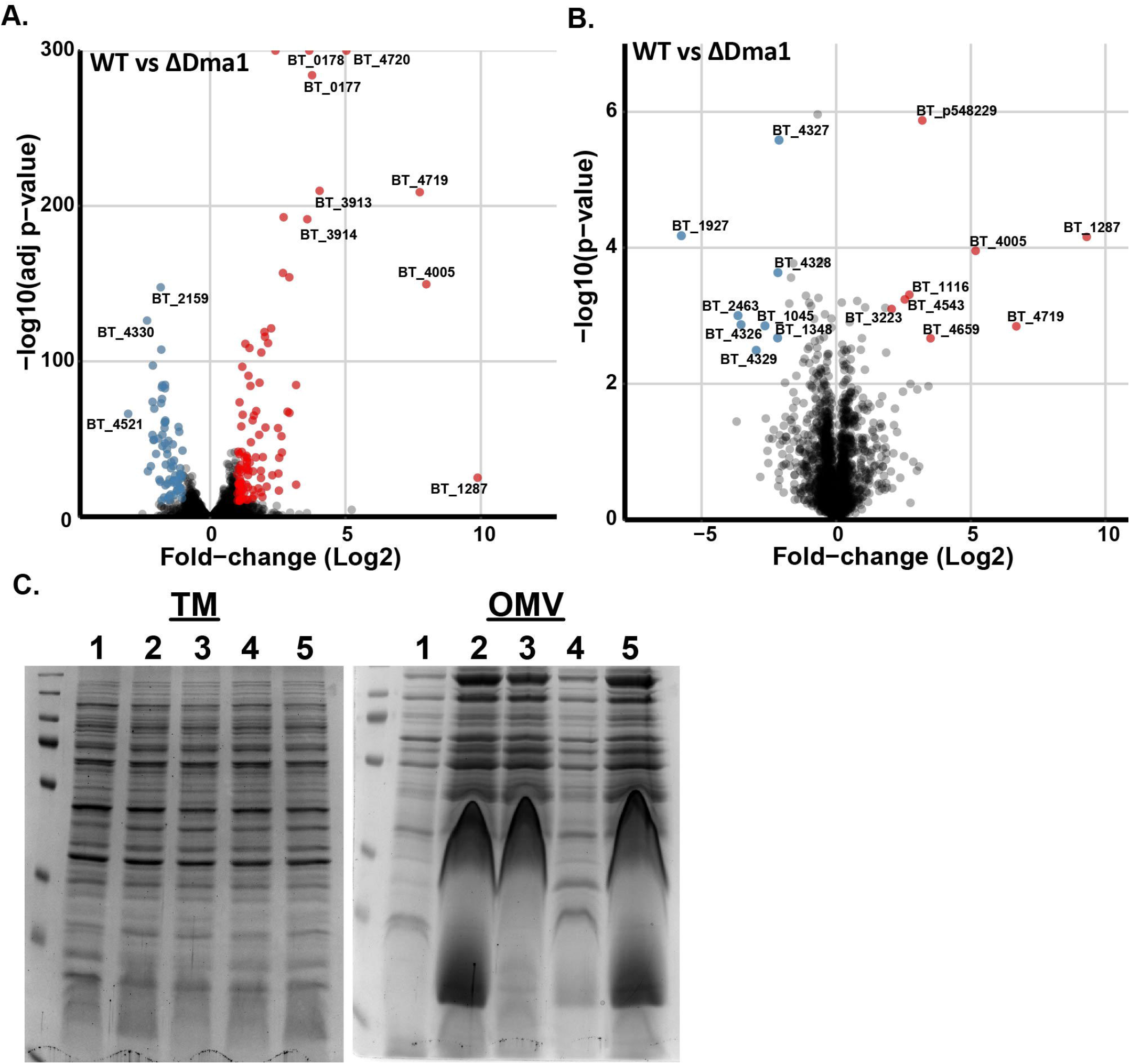
Member of the Dma1 regulon, NigD1 (BT_4005), is necessary to induce vesiculation. Volcano plot representations of (A) transcriptome and (B) proteome data comparing BT WT and BTΔDma1. (C) Coomassie Blue stain of TM and OMV fractions isolated from 1. WT, 2. ΔDma1, 3. Δ1287/Dma1, 4. ΔNigD1/Dma1, 5. ΔNigD2/Dma1. Left: Protein profile of mutant TMs are the same as WT, indicating that all changes are specific to the OMV fraction. Right: Deletion of NigD1 in the BTΔDma1 background restores the protein profile to WT. Samples were normalized by OD_600_ and run on 10% SDS-PAGE.

### Dma1 represents an unprecedented class of anti-sigma factor that spans both the IM and OM

Canonical anti-sigma factors contain a cytoplasmic domain, responsible for the sequestration of their cognate sigma-factors at the IM, connected via a transmembrane region to a periplasmic sensor domain (Otero-Asman et al. 2019). Dma1 is annotated as an OMP_b-brl_2 domain-containing protein. Structural modeling with Alphafold (Varadi et al. 2022) predicts that Dma1 contains an N-terminal alpha helical transmembrane region and a C-terminal eight stranded β-barrel domain connected via a long intrinsically disordered region **(Fig. 4A)**. Eight stranded β-barrel domain proteins are typically found in the OM of Gram-negative bacteria. In *Bf*, Reo, was shown to directly sequester its cognate sigma factor via its N-terminal region in the cytoplasm (Ndamukong et al. 2013). Interaction between Dma1 and Das1 is supported by our observation that deletion of Das1 reverts the phenotype of ΔDma1. Based on these observations, we propose an unprecedented domain architecture for Dma1, in which the C-terminal β-barrel is embedded in the OM; the N-terminal domain that interacts with the cognate sigma factor is in the cytoplasm anchored to the IM via an alpha helical transmembrane helix; and both domains are tethered via the intrinsically disordered region that traverses the periplasm **(Fig. 4B)**. Based on this, we propose that Dma1 is the founding member of the <u>D</u>ual <u>M</u>embrane-spanning <u>A</u>nti-sigma factors family. To provide support for this model, we performed Western blots of *Bt* cells co-expressing C-terminal His-tagged Dma1 and Das1. Cells were harvested during exponential and late stationary phase. Expression of Dma1 was enhanced when expressed alongside Das1. Wild-type *Bt* and cells expressing only Das1 were employed as controls. During exponential phase growth, a band corresponding to full-length (∼55kDa) and a C-terminal-containing fragment (∼20kDa) of Dma1 were detected **(Fig. 4C)**. At stationary phase, full-length Dma1 was no longer detected, with the concomitant appearance of an additional smaller fragment. These results suggest that Dma1 is temporally regulated **(Fig. 4C)**. Next, we generated cells expressing Dma1 tagged with both a C-terminal 10xHis tag and an N-terminal 3xFLAG tag. Whole cells (WCs), TMs, and OMVs were analyzed by Western blot. Full-length Dma1 (∼55kDa) was detected in the WC and TM fractions with both antibodies, confirming that Dma1 is translated and translocated as a complete and single polypeptide. Only C-terminal fragments were detected in the OMVs using the anti-His antibodies, indicating that, as predicted, the β-barrel domain is localized in the OM **(Fig. 4D and Fig. S9)**. In addition, our conclusion is supported by the MS analysis of wild-type OMVs, which only detected peptides containing the predicted C-terminal β-barrel **(Table S5)**. Taken together, our results provide strong evidence that Dma1 is an anti-sigma factor that spans both membranes, directly connecting the exterior of the cell with its cytoplasm.

**Figure 4:**
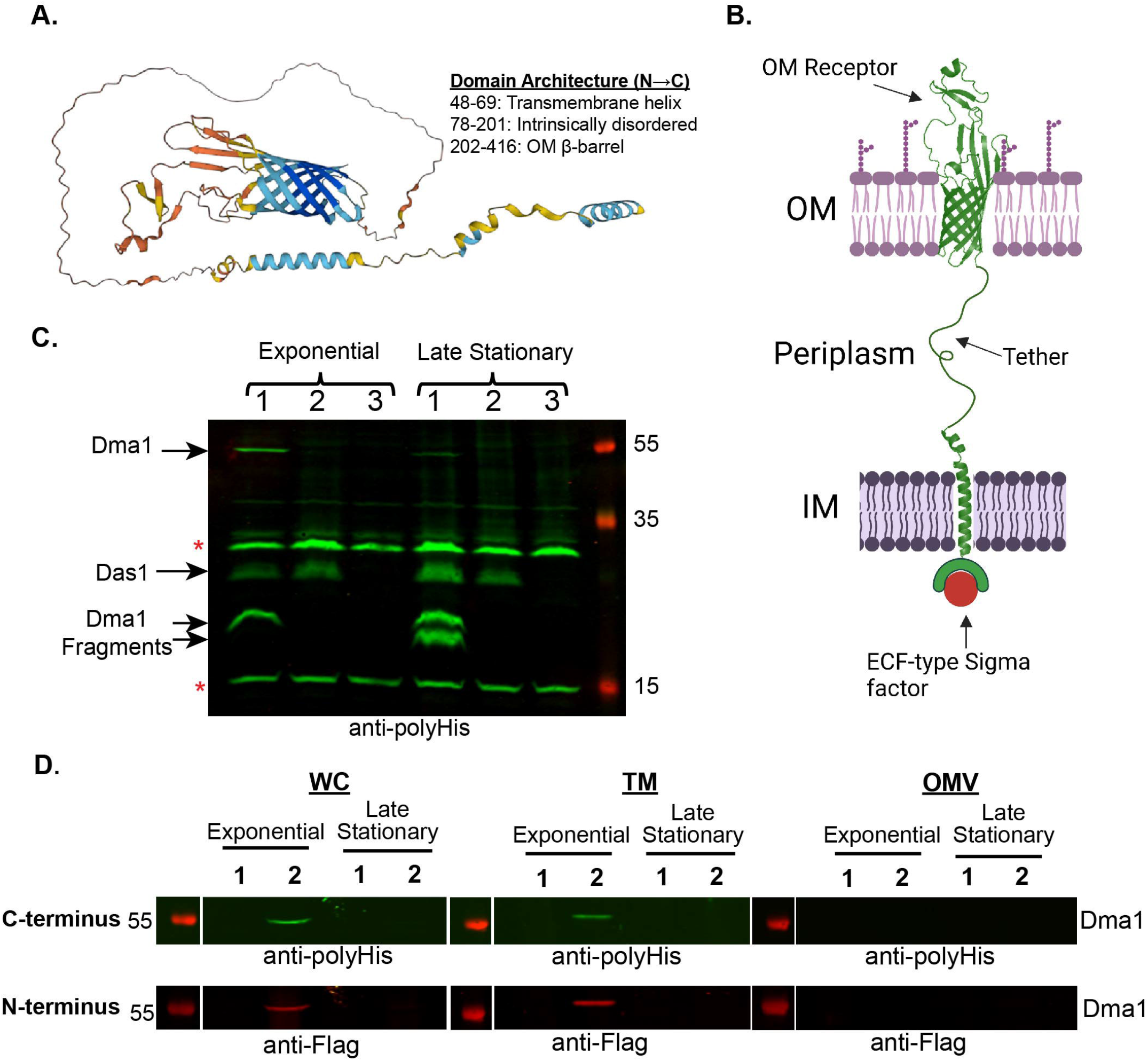
Dual Membrane-spanning 1 (Dma1) is the first protein shown to span both the inner and outer membrane of a Gram-negative bacterium. (A) AlphaFold predicted stucture of Dma1. (B) Proposed orientation of Dma1 in the membrane of *Bt* (Created with BioRender.com). (C) Western blot against anti-polyHis from WCs constitutively expressing 1. Dma1 and Das1 together, 2. Das1 alone, and 3. empty vector control. Bands adjacent to stars (*) are non-specific bands. Demonstrates that full-length Dma1 is present in WCs, and the state of the protein is growth-phase dependent. (D) Western blots of WCs, TMs, and OMVs collected from Bt strain constitutively expressing Dma1 containing a C-terminal His tag and an N-terminal Flag tag. Green channel is anti-polyHis, while the red channel is anti-Flag. Full-length Dma1 is present solely in the WC and TM fraction.

### Dual Membrane-spanning Anti-sigma factors are present throughout Bacteroidota

To determine whether proteins with similar domain architecture exist in *Bt*, we searched for sigma/anti-sigma pairs containing anti-sigma factors encoding a predicted β-barrel domain. We identified two additional proteins in *Bt*, Dma2 (BT_1558) and Dma3 (BT_2778), that are structurally similar to Dma1 **(Fig. 5A)**. Dma2 is encoded adjacent to a sigma factor, BT_1559 (Das2), while Dma3 is more complex, as it is predicted to contain the sigma factor fused to the rest of the polypeptide at the N-terminus **(Fig. 5A)**. A bioinformatic analyses showed that Dma1 is present in almost all Bacteroidota, while Dma2 and Dma3 are less prevalent. **Figure 5B** shows a phylogenetic analysis containing select members of the Bacteroidota representing various classes ranging from mammalian gut commensals to soil-dwelling microbes. Dma1 is present in almost all Bacteroidota chosen (27/29), while Dma2 (5/29) and Dma3 (10/29) were less prevalent **(Fig. 5B)**. Interestingly, Dma2 is only present in members of the genus *Bacteroides*, suggesting that this protein was recently acquired in the genus. Dma3 on the other hand is predominantly found in *Bacteroides* and *Prevotella* **(Fig. 5B and Fig. S10)**. We were unable to identify any structurally similar proteins in Gram-negative bacteria outside of Bacteroidota. To determine whether Dma2 and Dma3 also modulate OMV biogenesis in *Bt*, we generated clean deletion mutants in each gene and analyzed their protein profiles by SDS-PAGE. Mutation of Dma2 produced a phenotype similar to that of ΔDma1, which suggests that Dma2 also modulates OMV biogenesis. Deletion of its cognate sigma factor Das2 also restores WT levels of vesiculation **(Fig. S11)**. Mutation of Dma3 revealed no detectable phenotype by SDS-PAGE (data not shown).

**Figure 5:**
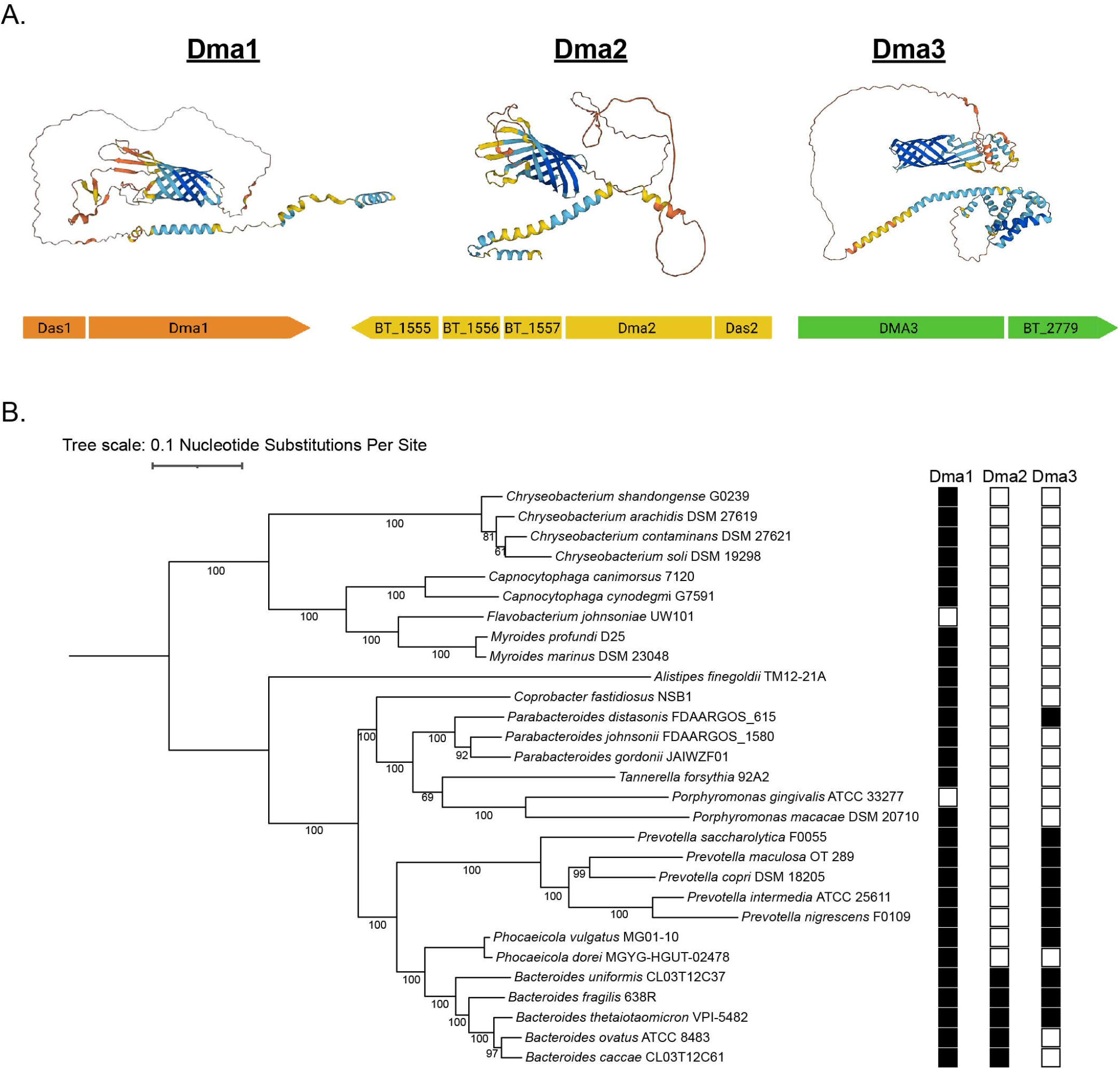
Members of the Dma family are present throughout Bacteroidota. (A) Genome analysis identified two additional proteins, Dma2 and Dma3, with structural similarity to Dma1. AlphaFold predicted structure of each Dma, along with their respective operons, are presented. (B) A maximum likelihood phylogenetic tree was constructed from twenty-nine core genomes of various Bacteroidota. Amino acid sequences of Dma1 (WP_008766767.1), Dma2 (WP_008762208.1), Dma3 (WP_011108473.1) from Bt served as references for identification in the other genomes.

## Discussion

Since their discovery in the 1960’s, many important roles have been proposed for OMVs (Bishop and Work 1965). However, to date very little is known about the mechanism(s) of bacterial vesiculation and its regulation. In this study, we aimed to advance our understanding of the mechanism of OMV biogenesis in Bacteroidota. To this aim, we developed a novel screening methodology to identify genes involved in OMV biogenesis. We discovered the Dma family, a novel family of anti-sigma factors with an unprecedented domain organization. Dma proteins span both membranes, possessing an extracellular and a cytoplasmic domain connected via a large, intrinsically disordered region that crosses the periplasm. We show that inactivation of Dma1 (BT_4721) or Dma2 (BT_1558) results in hypervesiculation in *Bt*.

We previously showed that labeling OM and OMV-specific proteins with distinct fluorescent markers is an effective way to visualize OMVs and distinguish them from byproducts of cell lysis (Sartorio et al., 2023). In this work, we labeled an OMV marker with NLuc instead of fluorescent proteins, which allowed us to quantify OMV production *in vitro* in a high throughput format **(Fig. 1C)**. Screens attempting to identify genes involved in OMV biogenesis have been performed in various bacteria (Kulp et. al., 2015); however, these were unable to establish causal associations between specific genes and OMV formation. We speculate that this is due to their use of non-specific markers, like LPS, OmpA, or phospholipids, as a readout for OMV production. Many of the genes identified in these studies led to membrane destabilization, which confounded the interpretation of the results, due to their inability to differentiate genuine OMVs from cell lysis. Although OMV cargo selection is not common in all bacteria, the methods developed here can be adapted to conduct OMV screens in other bacterial species.

Our model for regulation of OMV biogenesis in *Bt* is summarized in **Fig. 6**. We propose that Dma proteins sense extracellular stimuli or perturbations in the OM via their C-terminal β-barrel domain, which triggers a series of proteolytic events that liberates their cognate sigma factors to modulate gene expression and induce OMV production **(Fig. 6)**. By employing transcriptomic and proteomic analyses, we demonstrated that NigD1 is required for the induction of vesiculation in the absence of Dma1, upregulating the amount of LPS, proteins and lipids present in OMVs. NigD-like proteins are ubiquitous among Bacteroidota, but their functions have not been defined. NigD1 is encoded adjacent to genes required for biosynthesis and regulation of LPS and phospholipids. Our transcriptomic and proteomic analyses indicate that these genes/proteins are not differentially regulated between WT and ΔDma1. Bacteroidota membranes are rich in sphingolipids and other lipids not commonly found in other bacteria. More knowledge about regulation of LPS and lipid biosynthesis in *Bacteroides* will be required to fully understand how Dma1, Dma2 and NigD1 control OMV biogenesis. NigD2 and BT_1287 are part of the Dma1 regulon, but they are dispensable for the hypervesiculation phenotype. Therefore, it is possible that Dma1 regulate other processes in Bt.

**Figure 6:**
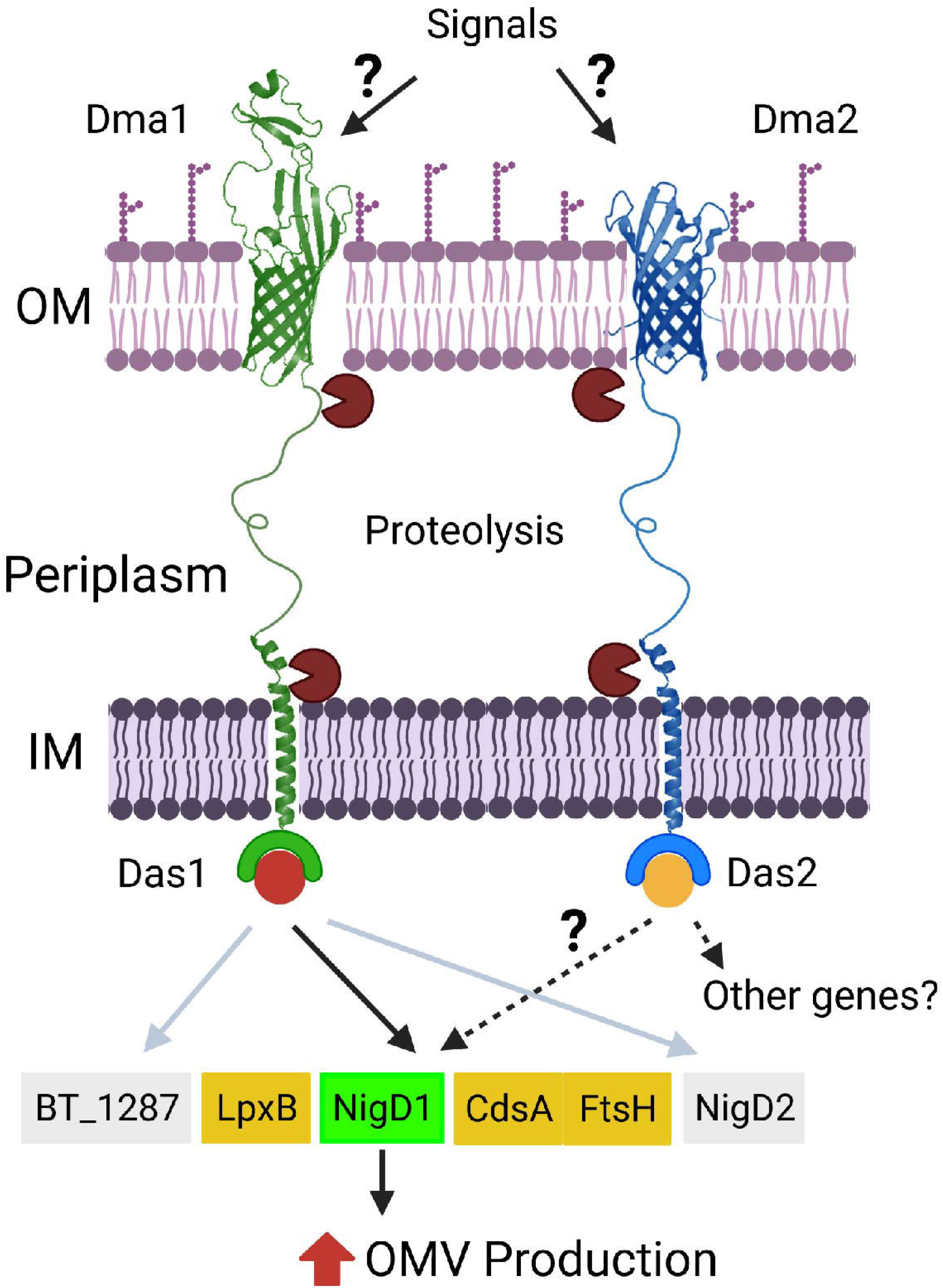
Schematic of Dma1- and Dma2-mediated induction of OMV biogenesis. In our model, Dma1 and Dma2 sense extracellular stimuli and/or perturbations in the OM. This leads to the proteolytic degradation of the proteins to release their cognate sigma factors, and subsequently modulate gene expression to induce OMV biogenesis. Dma1/Das1-mediated hypervesiculation is dependent upon NigD1, while the regulon of Dma2 is unclear.

Previous studies have investigated orthologs of Dma1. In *B. fragilis*, deletion of *Reo* was shown to increase fitness in response to oxidative stressors (Ndamukong et al. 2013). However, we found that Dma1 does not seem to play the same role in *Bt* **(Fig. S7)**. In *Phocaeicola vulgatus*, mutation of the Dma1 ortholog, M098_03760, conferred increased protection against the antimicrobial toxin BcpT. Authors hypothesize that this occurs by increasing LPS O-antigen length and preventing BcpT from binding lipid A core and destabilizing the OM (Evans et al. 2022). We did not analyze the structure of LPS in this work, but reports suggest that *Bt* lacks LPS, and instead makes lipooligosaccharide (LOS) (Jacobson et al. 2018). Therefore, Dma1 is not likely to have this function in *Bt*. Members of the Dma family likely play diverse roles in different bacterial species. In *Bt*, the Dma family impacts OMV biogenesis. Since OMVs have been implicated in stress response, it would be interesting to determine whether the fitness advantages observed in other studies are somehow related to increased vesiculation.

The unprecedented domain organization of the Dma family raises fundamental questions regarding their translocation and assembly. In Gram-negative bacteria, β-barrel <u>O</u>uter <u>M</u>embrane <u>P</u>roteins (OMPs) are trafficked to the periplasm by translocation through the Sec system, where chaperones, like surA, shuttle the unfolded OMPs to the <u>β</u>-barrel <u>A</u>ssembly <u>M</u>achine (BAM) for insertion into the OM (Knowles et al. 2009; Noinaj et al. 2017). Since Dma1 has domains inserted into both membranes, it is unclear whether Dma1 is localized via the BAM complex. It is tempting to speculate that Bacteroidota have evolved additional systems specifically to ensure the proper localization of the Dma proteins, but future studies are required to validate this hypothesis.

The presence of Dma1 proteolytic fragment provides clues about Dma1 regulation. In *E. coli* and similar organisms, there are typically two proteolytic events that occur at the inner and outer leaflet of the IM to inactivate anti-sigma factors (Kim and Young 2015; Otero-Asman et al. 2019). Two IM proteases, DegS and RseP, involved in RIP have been characterized (Li et al. 2009; Dartigalongue et al. 2001). Potential orthologs of these IM proteases are predicted to be present in *Bt*. Future work will be focused on understanding what are the signals that trigger Dma1 proteolysis, and the enzymes involved in the process.

Our screening led us to the discovery of the Dma family and their unusual domain organization. However, we also identified multiple mutants displaying hyper- and hypovesiculation phenotypes. We expect that the investigation of these mutants will further our understanding of the bacterial vesiculation mechanisms and will lead to identification of the hitherto elusive machinery responsible for OMV biogenesis.

## Acknowledgements

We thank all members of the Feldman lab for critically reading the manuscript. This work was supported by NIH grants R21AI168719 and R21AI151873 to M.F.F. N.E.S is supported by an Australian Research Council Future Fellowship (FT200100270) and an ARC Discovery Project Grant (DP210100362). We thank the Melbourne Mass Spectrometry and Proteomics Facility of The Bio21 Molecular Science and Biotechnology Institute for access to MS instrumentation.

## Supplementary Figures

**Figure S1: Hyper- and hypovesiculating strains were identified during OMV screening**. NanoGlow assays were performed using filtered supernatants isolated from each potential candidate. Total luminescent output was normalized to OD_600_. Candidates exhibiting a 1.5-fold increase in luminescence were considered hypervesiculating, while those displaying a 0.5-fold decrease were deemed hypovesiculating.

**Figure S2: ΔDma1 and ΔDas1 do not impact fitness in vitro**. Growth curves performed in BHI media for WT, ΔDas1, ΔDma1, ΔDas1-Dma1, and corresponding complemented strains.

**Figure S3: Protein composition of subcellular fractions is consistent between the wild-type and the mutants**. Principal component analysis (PCA) of WC (stars), TM (squares), and OMV (circles) proteomes from Bt grown in BHI media. Four biological replicates were performed for each condition.

**Figure S4: OMV cargo selection is not impacted by mutation of Dma1 operon**. Volcano plot representations of proteins enriched in the TM and OMV fractions. Integral membrane proteins are represented in blue, lipoproteins with LES motifs are indicated in red, lipoproteins lacking the LES motif are depicted in yellow, and soluble proteins are indicated in dark gray. (A) OMV cargo selection is maintained in BTΔDma1, indicating that these OMVs are not the result of cell lysis. (B) OMV cargo selection is maintained in BTΔDas1 and BTΔDas1-Dma1.

**Figure S5: Hypervesiculation causes ΔDma1 to secrete significantly more membrane lipids**. Total lipids were isolated from (A) TM and (B) OMV fractions from WT, ΔDma1, and ΔDma1_Comp_. Lipids were relativized to an 18:0-20:4 phosphoinositol internal standard (IS). Red squares denote sphingolipids, dihydroceramides (DHC), ethanolamine-phosphoceramides (EPC), inositolphosphoceramide (IPC). Blue squares are phospholipids, phosphatidylethanolamine (PE), phosphatidylserine (PS), and phosphatidylinositol (PI). Yellow square represent amino lipids, glycylserine dipeptide lipids (GS), glycylserine phosphoryl diacylglycerol (GS-PA), N-(3-O-Acyl)acyl glycylserine phosphoryl dihydroceramide (GS-PDHC).

**Figure S6: Amino acid sequence alignment of Dma1 and *Bf* Reo**. Dma1 show 56% sequence identity with Reo. Amino acids highlighted in red represent high consensus levels (90%), while those in blue represent low consensus (50%). The predicted anti-sigma domain is shown in brackets. Alignment was done using MultAlin version 5.4.1.

**Figure S7: ECF21 family sigma factor, Das1, is required for hypervesiculation**. Coomassie Blue stain comparing protein profiles between WT, ΔDas1, ΔDma1, and ΔDas1-Dma1. This shows that deletion of Das1 in the ΔDma1 background restores WT levels of vesiculation. Samples were normalized by OD_600_ values and run on 10% SDS-PAGE gel.

**Figure S8: Mutants in ΔDas1 and ΔDma1 are not attenuated in aerobic stress**. Aerobic exposure stress tests comparing WT, ΔDas1, ΔDma1, and their corresponding complemented strains incubated in air for different times. Strains were cultured overnight, then diluted to the equivalent of OD=0.1 for the initial spot. Each subsequent spot is a ten-fold dilution of the previous one.

**Figure S9: Dual Membrane-spanning 1 (Dma1) is the first protein shown to span both the inner and outer membrane of a Gram-negative bacterium**. Western blots of WCs, TMs, and OMVs collected from Bt strain constitutively expressing Dma1 containing a C-terminal His tag and an N-terminal Flag tag. Green channel is anti-polyHis, while the red channel is anti-Flag. Full-length Dma1 is present solely in the WC and TM fraction.

**Figure S10: Schematic of gene synteny of *Bt* Dma1-3**. Gene synteny for different Bacteroidota were compared to (A) Dma1, (B) Dma2, and (C) Dma3 from Bt. Genes of the same color are conserved in different species, while those in grey differ.

**Figure S11: ΔDma2 induces OMV biogenesis in a similar manner to ΔDma1**. Coomassie Blue stain comparing protein profiles between WT, ΔDma2, ΔDas2, and ΔDma2-Das2. This gel shows that mutation of Dma2 induces vesiculation, and this phenotype is dependent on its cognate sigma factor, Das2. Samples were normalized by OD600 values and run on 10% SDS-PAGE gel.

