## Supplemental Tables for "Dual Membrane-spanning Anti-Sigma Factors Regulate Vesiculation in Gut Bacteroidota"

### Table S1: Strain, Plasmids, and Oligonucleotides

| **Strains used in this study** | | | |
|---|---|---|---|
| **Name** | **Strain** | **Features** | **Reference/Source** |
| *Escherichia coli* | s17-1λ*pir* | Conjugation donor strain to introduce plasmids into *B. thetaiotaomicron* | Jeffrey I. Gordon Laboratory |
| *B. thetaiotaomicron* | VPI-5482 | Wild-type strain. $Erm^S$ | Jeffrey I. Gordon Laboratory |
| *B. thetaiotaomicron* | VPI-5482 pwwBoINL-Nluc | Expressing *B. ovatus* inulinase fused to Nluc | This study |
| *B. thetaiotaomicron* | VPI-5482 Δ*das1* | *das1* (*bt_4720*) deletion mutant | This study |
| *B. thetaiotaomicron* | VPI-5482 Δdma1 | *dma1* (*bt_4721*) deletion mutant | This study |
| *B. thetaiotaomicron* | VPI-5482 Δ*das1-dma1* | *das1* (*bt_4720*)-*dma1* (*bt_4721*) double deletion mutant | This study |
| *B. thetaiotaomicron* | VPI-5482 Δ*das2* | *das2* (*bt_1559*) deletion mutant | This study |
| *B. thetaiotaomicron* | VPI-5482 Δ*dma2* | *dma2* (*bt_1558*) deletion mutant | This study |
| *B. thetaiotaomicron* | VPI-5482 Δ*das2-dma2* | *das2* (*bt_1559*)-*dma2* (*bt_1558*) double deletion mutant | This study |
| *B. thetaiotaomicron* | VPI-5482 Δ*das1* | *das1* (*bt_4720*) deletion mutant, complemented | This study |
| *B. thetaiotaomicron* | VPI-5482 Δ*dma1* pwwDma1-His | *dma1* (*bt_4721*) deletion mutant, complemented | This study |
| *B. thetaiotaomicron* | VPI-5482 Δ*das1-dma1* pwwDas1-Dma1 | *das1* (*bt_4720*)-*dma1* (*bt_4721*) double deletion mutant, complemented | This study |
| *B. thetaiotaomicron* | VPI-5482 Δ*dma1* pwwFLAG-Dma1-His | *dma1* (*bt_4721*) deletion mutant expressing pwwFLAG-Dma1-His | This study |
| **Plasmids used in this study** | | | |
| **Name** | **Resistance** | **Features** | **Reference/Source** |
| pSAM-Bt | $Amp^R$,$Erm^R$ | Vector base to perform transposon mutagenesis | Goodman et. al., 2009 |
| pSAM-Bt_Tet | $Amp^R$,$Tet^R$ | pSAM-Bt backbone containing tetracycline resistance cassette | This work |
| pSIE1 | $Amp^R$,$Erm^R$,$aTc^S$ | P1T_DP-GH023_*ss-bte1*; P1T_DP-GH023_*ss-bfe1*; $P_{BFP1E6}$_tetR | Bencivenga-Barry et. al., 2020 |
| pSIE1_das1 | $Amp^R$,$Erm^R$,$aTc^S$ | pSIE1 deletion construct for *B. thetaiotaomicron* VPI-5482 *das1* | This study |
| pSIE1_dma1 | $Amp^R$,$Erm^R$,$aTc^S$ | pSIE1 deletion construct for *B. thetaiotaomicron* VPI-5482 *dma1* | This study |
| pSIE1_das1-dma1 | $Amp^R$,$Erm^R$,$aTc^S$ | pSIE1 deletion construct for *B. thetaiotaomicron* VPI-5482 *das1*& *dma1* | This study |
| pSIE1_das2 | $Amp^R$,$Erm^R$,$aTc^S$ | pSIE1 deletion construct for *B. thetaiotaomicron* VPI-5482 *das2* | This study |
| pSIE1_dma2 | $Amp^R$,$Erm^R$,$aTc^S$ | pSIE1 deletion construct for *B. thetaiotaomicron* VPI-5482 *dma2* | This study |
| pSIE1_das2-dma2 | $Amp^R$,$Erm^R$,$aTc^S$ | pSIE1 deletion construct for *B. thetaiotaomicron* VPI-5482 *das2* & *dma2* | This study |

| | | | |
|---|---|---|---|
| pSIE1_BT1287 | $Amp^R$,$Erm^R$,$aTc^S$ | pSIE1 deletion construct for *B. thetaiotaomicron* VPI-5482 BT_1287 | This study |
| pSIE1_nigD1 | $Amp^R$,$Erm^R$,$aTc^S$ | pSIE1 deletion construct for *B. thetaiotaomicron* VPI-5482 *NigD1* (BT_4005) | This study |
| pSIE1_nigD2 | $Amp^R$,$Erm^R$,$aTc^S$ | pSIE1 deletion construct for *B. thetaiotaomicron* VPI-5482 *NigD2* (BT_4719) | This study |
| pWW3452 | $Amp^R$,$Erm^R$ | AmpR-ermG-RP4/R6K-[P_BfP1E6-RBSlp-LPGFP- tag-Term]-NBU2, AmpR ErmR | Whitaker et. al., 2017 |
| pWW3867 | $Amp^R$,$Erm^R$ | AmpR-ermG-RP4/R6K-[P_BT1311-RBSphageGFP-tag-Term]-NBU2, AmpR ErmR | Whitaker et. al., 2017 |
| pwwBoINL-Nluc | $Amp^R$,$Erm^R$ | pWW3867containing Inulinase-Nluc fusion | This study |
| pwwNLuc | $Amp^R$,$Erm^R$ | pWW3867containing Nluc | This study |
| pwwEV | $Amp^R$,$Erm^R$ | pWW3867 backbone | Whitaker et. al., 2017 |
| pwwDas1-His | $Amp^R$,$Erm^R$ | pWW3867 containing Das1 with C-terminal 6xHis tag | This study |
| pwwDma1-His | $Amp^R$,$Erm^R$ | pWW3867 containing Dma1 with C-terminal 10xHis tag | This study |
| pwwFLAG-Dma1-His | $Amp^R$,$Erm^R$ | pWW3867 containing Dma1 with an N-terminal 3xFlag tag and a C-terminal 10xHis tag | This study |
| pwwDas1-Dma1-His | $Amp^R$,$Erm^R$ | pWW3867 containing Das1 and Dma1 with C-terminal His tags | This study |

**Oligonucleotides used in this study**

| Name | Sequence | Template | Description |
|---|---|---|---|
| F_Nluc | GTCTTCACACTCGAAGATTTC | pLenti6.2-Nanoluc-ccdB vector | For cloning into pWW3867 |
| R_RpoD_Nluc(6His) | TCGAGCTAATCAGCTAGGATTTAG TGATGATGATGATGATGACCCGCC AGAATGCGTTC | pLenti6.2-Nanoluc-ccdB vector | For cloning into pWW3867 |
| F_BoIN-RpoD | TCCAAATCTGTTTTTAAAGAATGAA GATAAATAAATTCTTAATAAGCGG | *B. ovatus* ATCC 8483 genomic DNA | For cloning into pWW3867 |
| R_BoIN(5F)Nluc | AAATCTTCGAGTGTGAAGACAGAT CCTCCTCCTCCTTTCTTAGCGCTT AGATAATG | *B. ovatus* ATCC 8483 genomic DNA | For cloning into pWW3867 |
| F_pww-NLuc-His linear | GTCTTCACACTCGAAGATTTC | pLenti6.2-Nanoluc-ccdB vector | For cloning into pWW3867 |
| R_pww-NLuc | TCGAGCTAATCAGCTAGGATTTAG TGATGATGATGATGATGACCCGCC AGAATGCG | pLenti6.2-Nanoluc-ccdB vector | For cloning into pWW3867 |
| F_BT4720-Downstream | GCGGCCGCTCTAGAACTAGTCGC CTTCGTCGTTATG | *B. thetaiotaomicron* VPI-5482 genomic DNA | For cloning into pSIE1 |
| R_BT4720-Downstream | TTTCTCTTCCATATCTCTACTTGCT TATACACGTGTTTACC | *B. thetaiotaomicron* VPI-5482 genomic DNA | For cloning into pSIE1 |
| F_BT4720-Upstream | GTAAACACGTGTATAAGCAAGTAG AGATATGGAAGAGAAAGAATTATG | *B. thetaiotaomicron* VPI-5482 genomic DNA | For cloning into pSIE1 |
| R_BT4720-Upstream | GATTAGCATTATGAGGATCCATTG CTGATCATTGGGTATG | *B. thetaiotaomicron* VPI-5482 genomic DNA | For cloning into pSIE1 |
| F_BT4721_Up | GCGGCCGCTCTAGAACTAGTTGG TCAGGATCTCTTTTATTTCTCTTAC CT | *B. thetaiotaomicron* VPI-5482 genomic DNA | For cloning into pSIE1 |

| | | | |
|---|---|---|---|
| R_BT4721_Up | ATCTCTACCTATCCGTTATTCATTATCCATTC | *B. thetaiotaomicron* VPI-5482 genomic DNA | For cloning into pSIE1 |
| F_BT4721_Down | AATAACGGATAGGTAGAGATTAGAACCCGGTGTTGCTTATTTCTTTGATGATG | *B. thetaiotaomicron* VPI-5482 genomic DNA | For cloning into pSIE1 |
| R_BT4721_Down | AAGATTAGCATTATGAGGATCCGGTGACGTATGTTTCAGTGTTC | *B. thetaiotaomicron* VPI-5482 genomic DNA | For cloning into pSIE1 |
| R_BT4720/21-Downstream | ATATAGCTCAACTATATTGATTGCTTATACACGTGTTTACC | *B. thetaiotaomicron* VPI-5482 genomic DNA | For cloning into pSIE1 |
| F_BT4721/20-Upstream | GTAAACACGTGTATAAGCAATCAATATAGTTGAGCTATATATTTTAAAAGC | *B. thetaiotaomicron* VPI-5482 genomic DNA | For cloning into pSIE1 |
| F_BT1558 Downsteam | GCGGCCGCTCTAGAACTAGTCAGGCACAGTGTCATAG | *B. thetaiotaomicron* VPI-5482 genomic DNA | For cloning into pSIE1 |
| R_BT1558 Downstream | CGCTGATGCCGGGATGACTGTCCTTATAAAAATGAAAAAAACACTTATG | *B. thetaiotaomicron* VPI-5482 genomic DNA | For cloning into pSIE1 |
| F_BT1558 Upstream | TTTTTTCATTTTTATAAGGACAGTCATCCCGGC | *B. thetaiotaomicron* VPI-5482 genomic DNA | For cloning into pSIE1 |
| R_BT1558 Upstream | GATTAGCATTATGAGGATCCCGTGCTGCAGCTG | *B. thetaiotaomicron* VPI-5482 genomic DNA | For cloning into pSIE1 |
| F_BT1559 Downstream | GCGGCCGCTCTAGAACTAGTGCTTTCAGCGGGATG | *B. thetaiotaomicron* VPI-5482 genomic DNA | For cloning into pSIE1 |
| R_BT1559 Downstream | ACACCCTCTATGCCAAACAATGAAATAACCGATCTGTTTCG | *B. thetaiotaomicron* VPI-5482 genomic DNA | For cloning into pSIE1 |
| F_BT1559 Upstream | GAAACAGATCGGTTATTTCATTGTTTGGCATAGAGGG | *B. thetaiotaomicron* VPI-5482 genomic DNA | For cloning into pSIE1 |
| R_BT1559 Upstream | GATTAGCATTATGAGGATCCTTGGATACCCGCAAG | *B. thetaiotaomicron* VPI-5482 genomic DNA | For cloning into pSIE1 |
| F_BT1287_Dwn | GCGGCCGCTCTAGAACTAGTCCGGTCATCCTTTACG | *B. thetaiotaomicron* VPI-5482 genomic DNA | For cloning into pSIE1 |
| R_BT1287_Dwn | TAAATAATTAAACAGTCACCTTTAGTTAAAGTTTTCATACTTGTTAAAC | *B. thetaiotaomicron* VPI-5482 genomic DNA | For cloning into pSIE1 |
| F_BT1287_Up | GTATGAAAACTTTAACTAAAGGTGACTGTTTAATTATTTACAG | *B. thetaiotaomicron* VPI-5482 genomic DNA | For cloning into pSIE1 |
| R_BT1287_Up | GATTAGCATTATGAGGATCCAGAGGCGAATGCG | *B. thetaiotaomicron* VPI-5482 genomic DNA | For cloning into pSIE1 |
| F_BT4005_Dwn | GCGGCCGCTCTAGAACTAGTTAGGAAATCCTACGGTAG | *B. thetaiotaomicron* VPI-5482 genomic DNA | For cloning into pSIE1 |
| R_BT4005_Dwn | AGTTATTTATAAAAGTTTTGATTTGTTCCTCTCTTTTTAATTC | *B. thetaiotaomicron* VPI-5482 genomic DNA | For cloning into pSIE1 |
| F_BT4005_Up | TTAAAAAGAGAGGAACAAATCAAAACTTTTATAAATAACTAAACCTAAAC | *B. thetaiotaomicron* VPI-5482 genomic DNA | For cloning into pSIE1 |
| R_BT4005_Up | GATTAGCATTATGAGGATCCACCTGTTCCTCGC | *B. thetaiotaomicron* VPI-5482 genomic DNA | For cloning into pSIE1 |
| F_BT4719_Dwn | GCGGCCGCTCTAGAACTAGTGAAGTGCTGTTTTTGCC | *B. thetaiotaomicron* VPI-5482 genomic DNA | For cloning into pSIE1 |
| R_BT4719_Dwn | AAAAAACATTTTTATAGTATATCTAGTAATTATAGAAACACTC | *B. thetaiotaomicron* VPI-5482 genomic DNA | For cloning into pSIE1 |
| F_BT4719_Up | TGTTTCTATAATTACTAGATATACTATAAAAATGTTTTTTAATGAGTACTATTTAC | *B. thetaiotaomicron* VPI-5482 genomic DNA | For cloning into pSIE1 |
| R_BT4719_Up | GATTAGCATTATGAGGATCCTCTTTCTCTTCCATATCTCTAC | *B. thetaiotaomicron* VPI-5482 genomic DNA | For cloning into pSIE1 |
| F_RpoD_BT4720-His | TCCAAATCTGTTTTTAAAGAATGGAAGAATTTGAGTTGTCG | *B. thetaiotaomicron* VPI-5482 genomic DNA | For cloning into pWW3867 |

| | | | |
|---|---|---|---|
| R_RpoD_BT4720-His | TCGAGCTAATCAGCTAGGATCTAG TGATGATGATGATGATGACCTCCG TTATTCATTATCCATTCTTTTAC | *B. thetaiotaomicron* VPI-5482 genomic DNA | For cloning into pWW3867 |
| F_pwwRpoD_BT4721-His | CCAAATCTGTTTTTAAAGAATGGA AGAGAAAGAATTATGG | *B. thetaiotaomicron* VPI-5482 genomic DNA | For cloning into pWW3867 |
| R_pwwRpoD_BT4721-His | TCGAGCTAATCAGCTAGGATCTAA TGATGATGATGGTGATGATGATGA TGATGACCATAAGTAAGTCGGATA CC | *B. thetaiotaomicron* VPI-5482 genomic DNA | For cloning into pWW3867 |
| R_RpoD_BT4720/21-His | TTTCTCTTCCATATCTCTACCTAGT GATGATGATGATGATGACCTCCGT TATTCATTATCCATTCTTTTAC | *B. thetaiotaomicron* VPI-5482 genomic DNA | For cloning into pWW3867 |
| F_pwwRpoD_BT4721/20-His | CATCATCACTAGGTAGAGATATGG AAGAGAAAGAATTATGG | *B. thetaiotaomicron* VPI-5482 genomic DNA | For cloning into pWW3867 |
| F_FLAG-4721_lin | ATGGATTACAAGGACGATG | pWW3452 DNA | For cloning into pWW3452 |
| R_N-term_3x-FLAG_BT4721 | ATCCATAATTCTTTCTCTTCTCCAG ATTTATCATCATCG | pWW3452 DNA | For cloning into pWW3452 |
| F_Phg_FLAG-BT4721-His | ACGATGATGATAAATCTGGAGAAG AGAAAGAATTATGGATG | *B. thetaiotaomicron* VPI-5482 genomic DNA | For cloning into pWW3452 |
| R_pwwRpoD_BT4721-His | TCGAGCTAATCAGCTAGGATCTAA TGATGATGATGGTGATGATGATGA TGATGACCATAAGTAAGTCGGATA CC | *B. thetaiotaomicron* VPI-5482 genomic DNA | For cloning into pWW3452 |

| Gene Interrupted | Putative Function | Phenotype | Number of Hits |
|---|---|---|---|
| BT_4721 | OMP_b-brl_2 domain containing protein | + | 4 |
| BT_4261 | DUF4847 domain-containing protein | - | 2 |
| BT_3709 | Elongation factor P | + | 3 |
| BT_3422 | MG2 domain protein | + | 1 |
| BT_3341 | SbsA Ig-like domain-containing protein | - | 1 |
| BT_2493 | ROK family transcriptional repressor | + | 4 |
| BT_2235 | Unknown | - | 1 |
| BT_1115 | Aldo/keto reductase | + | 1 |
| BT_0753 | Anti-sigma factor | - | 1 |
| BT_0372 | Aldose 1-epimerase | + | 1 |

**Table S2: List of candidate genes identified during OMV Screen.** Hyper- and hypovesiculating mutants were confirmed by isolating TM and OMV fractions from the wild-type and transposon mutants, performing western blots using anti-polyHis antibodies, and quantifying the relative amount of INL-Nluc in the OMV fraction by Li-cor. Hypervesiculating strains are those denoted with a plus (+) because they exhibited a 1.5-fold increase in signal, while hypovesiculating strains are those with a minus (-) and displayed a 0.5-fold decrease in signal.

**Table S3: RNA Sequencing-ΔDma1 Top 25 Most Upregulated Genes**

| Gene (New Locus Tag) | Gene (Old Locus Tag) | $Log_2FC$ | Padj | Proposed Function |
|---|---|---|---|---|
| BT_RS06495 | BT_1287 | 9.879633157 | 7.34E-26 | DUF4840 domain-containing protein |
| BT_RS20210 | BT_4005 | 7.986054337 | 2.34E-150 | NigD-like protein |
| BT_RS23775 | BT_4719 | 7.743518851 | 1.26E-209 | NigD-like protein |
| BT_RS16205 | NO | 5.229630823 | 3.24E-05 | Smalltalk protein |
| BT_RS23780 | BT_4720 | 5.034076276 | 0 | Sigma-70 family RNA polymerase sigma factor |
| BT_RS19750 | BT_3913 | 4.044393237 | 1.72E-210 | DUF5034 domain-containing protein |
| BT_RS00870 | BT_0177 | 3.767930416 | 6.95E-285 | NigD-like protein |
| BT_RS00875 | BT_0178 | 3.662030315 | 0 | YIP1 family protein |
| BT_RS19755 | BT_3914 | 3.590716188 | 3.32E-192 | Hypothetical protein |
| BT_RS11715 | BT_2315 | 3.579969558 | 0.0040536 | Hypothetical protein |
| BT_RS24670 | NO | 3.287601612 | 0.02828646 | Hypothetical protein |
| BT_RS05220 | BT_1038 | 3.178373436 | 2.30E-21 | endo-beta-N-acetylglucosaminidase family protein |
| BT_RS20385 | BT_4039 | 3.177706063 | 1.72E-85 | TonB-dependent receptor |
| BT_RS23805 | BT_4725 | 2.955721786 | 0.00240863 | RagB/SusD family nutrient uptake outer membrane protein |
| BT_RS05230 | BT_1040 | 2.931638602 | 1.49E-67 | SusC/RagA family TonB-linked outer membrane protein |
| BT_RS22810 | BT_4523 | 2.926352615 | 7.39E-155 | Type I restriction endonuclease EcoR124II |
| BT_RS14080 | BT_2779 | 2.860410832 | 2.41E-68 | Class I SAM-dependent methyltransferase |
| BT_RS10570 | BT_2086 | 2.691908858 | 1.37E-157 | Linear amide C-N hydrolase |
| BT_RS05210 | BT_1036 | 2.653696133 | 4.33E-42 | DUF1735 domain-containing protein |
| BT_RS22805 | BT_4522 | 2.637320216 | 1.54E-52 | Type I restriction endonuclease |
| BT_RS05225 | BT_1039 | 2.5531964 | 1.13E-38 | SusD/RagB family nutrient-binding outer membrane lipoprotein |
| BT_RS22645 | BT_4490 | 2.541660806 | 1.52E-17 | DUF5025 domain-containing protein |
| BT_RS05215 | BT_1037 | 2.524779245 | 1.45E-28 | DUF1735 and LamG domain-containing protein |
| BT_RS12945 | BT_2560 | 2.50963332 | 8.48E-58 | TonB-dependent receptor |
| BT_RS14845 | NO | 2.460763495 | 9.28E-08 | Smalltalk protein |

**Table S3: RNA Sequencing-ΔDma1 Top 25 Most Downregulated Genes**

| Gene (New Locus Tag) | Gene (Old Locus Tag) | $Log_2FC$ | Padj | Proposed Function |
|---|---|---|---|---|
| BT_RS22800 | BT_4521 | -3.0243084 | 5.47E-67 | Tyrosine-type recombinase/integrase |
| BT_RS15310 | BT_3017 | -2.497832498 | 0.01664662 | Acid phosphatase |
| BT_RS21860 | BT_4330 | -2.326456539 | 5.53E-127 | Nucleoside permease |
| BT_RS09755 | BT_1926 | -2.295427189 | 5.78E-30 | OmpA/MotB domain protein |
| BT_RS18675 | BT_3704 | -2.184786525 | 3.07E-33 | Glycoside hydrolase family 13 protein |
| BT_RS04750 | BT_0943 | -2.127262731 | 2.26E-53 | Penicillin-binding protein 2B (PBP-2B) |
| BT_RS13245 | BT_2619 | -2.123420723 | 1.05E-74 | Histidine kinase |
| BT_RS21840 | BT_4326 | -2.112073399 | 4.79E-98 | ATP-binding cassette domain-containing protein |
| BT_RS16435 | BT_3244 | -2.081737712 | 3.53E-50 | BACON domain-containing protein |
| BT_RS04760 | BT_0945 | -2.078233418 | 3.80E-50 | Hypothetical protein |
| BT_RS14350 | BT_2830 | -2.060239218 | 2.68E-70 | HU family DNA-binding protein |
| BT_RS09760 | BT_1927 | -1.993238565 | 3.54E-41 | Hypothetical protein |
| BT_RS16430 | BT_3243 | -1.909968855 | 4.73E-51 | DUF4302 domain-containing protein |
| BT_RS13240 | BT_2618 | -1.881918801 | 9.75E-09 | Two-component system response regulator |
| BT_RS20375 | BT_4037 | -1.872169035 | 1.38E-24 | Hypothetical protein |
| BT_RS15060 | BT_2972 | -1.835815403 | 2.91E-23 | Class I SAM-dependent methyltransferase |
| BT_RS10930 | BT_2159 | -1.822844005 | 2.25E-148 | Gfo/Idh/MocA family oxidoreductase |
| BT_RS21835 | BT_4325 | -1.809901243 | 8.80E-74 | ABC transporter permease |
| BT_RS08875 | BT_1751 | -1.805310189 | 2.07E-07 | Glycine betaine/L-proline ABC transporter ATP-binding protein |
| BT_RS21845 | BT_4327 | -1.801115206 | 2.77E-108 | DUF4836 family protein |
| BT_RS13230 | BT_2615 | -1.793091582 | 1.50E-07 | Group II intron reverse transcriptase/maturase |
| BT_RS04755 | BT_0944 | -1.791690553 | 3.24E-11 | Hypothetical protein |
| BT_RS13260 | BT_2622 | -1.780022703 | 5.06E-84 | Alpha-glucuronidase |
| BT_RS21850 | BT_4328 | -1.774306654 | 1.52E-76 | 16S rRNA (uracil(1498)-N(3))-methyltransferase |
| BT_RS10860 | BT_2145 | -1.764647845 | 2.52E-85 | Adenosylcobalamin-dependent ribonucleoside-diphosphate reductase |

**Table S4: Proteomics- ΔDma1 Top 25 Most Upregulated Genes**

| Protein ID | Gene | Fold Change | -Log(P-value) | Proposed Function |
|---|---|---|---|---|
| Q8A887 | BT_1287 | 9.30394 | 4.16088 | DUF4840 domain-containing protein |
| Q89YL2 | BT_4719 | 6.68855 | 2.84572 | NigD-like protein |
| Q8A0L6 | BT_4005 | 5.17626 | 3.95591 | NigD-like protein |
| Q89YS2 | BT_4659 | 3.50177 | 2.6692 | SusD homolog |
| Q8AB86 | BT_0224 | 3.42438 | 1.96457 | Lipocalin-like protein |
| Q8ABW9 | BT_p548229 | 3.19438 | 5.8735 | Fimbrillin family protein |
| Q8A2W6 | BT_3189 | 3.012 | 1.91681 | Unknown |
| Q8ABX1 | BT_p548227 | 2.75674 | 1.99822 | DUF3575 domain-containing protein |
| Q8A8Q1 | BT_1116 | 2.7145 | 3.31 | PPC domain-containing protein |
| Q89Z34 | BT_4543 | 2.54599 | 3.24113 | Putative type I restriction enzyme specificity protein |
| Q8A174 | BT_3792 | 2.47739 | 1.88394 | alpha-1,6-mannanase |
| Q8A0W2 | BT_3909 | 2.1813 | 1.63533 | Unknown |
| Q8A2T3 | BT_3222 | 2.146 | 1.39245 | DUF4848 domain-containing protein |
| Q8A2T2 | BT_3223 | 2.05685 | 3.09966 | OMP_b-brl domain-containing protein |
| Q8A6W1 | BT_1765 | 1.85376 | 2.45838 | Levanase (2,6-beta-D-fructofuranosidase) |
| Q8A3A2 | BT_3052 | 1.82297 | 3.1166 | AraC family transcriptional regulator |
| Q8A8I7 | BT_1180 | 1.70579 | 1.8163 | Glycoside transferase family 4 |
| Q8A0E0 | BT_4081 | 1.69799 | 1.36892 | SusC homolog |
| Q8A113 | BT_3858 | 1.64563 | 1.41392 | Alpha-1,2-mannosidase |
| Q8A9K1 | BT_0814 | 1.60821 | 1.70922 | BamA/TamA family outer membrane protein |
| Q8AB18 | BT_0294 | 1.60238 | 1.35371 | Carboxypeptidase regulatory-like domain-containing protein |
| Q8A5R5 | BT_2173 | 1.38299 | 1.42681 | SusD homolog |
| Q8A151 | BT_3816 | 1.33909 | 1.3211 | Penicillin-binding protein 2 (PBP-2) |
| Q8AA46 | BT_0619 | 1.27055 | 1.6609 | Ion-translocating oxidoreductase complex subunit D, rnfD |
| Q8A109 | BT_3862 | 1.18718 | 1.70627 | Endo-alpha-mannosidase |

**Table S4: Proteomics- ΔDma1 Top 25 Most Downregulated Genes**

| Protein ID | Gene | Fold Change | -Log(P-value) | Proposed Function |
|---|---|---|---|---|
| Q8A6F7 | BT_1927 | -5.75443 | 4.17785 | Unknown |
| Q8ABI4 | BT_0126 | -3.70228 | 1.44507 | Six-hairpin glycosidase |
| Q8A4Y4 | BT_2463 | -3.65105 | 3.00365 | RNA polymerase ECF-type sigma factor |
| Q89ZQ0 | BT_4326 | -3.53782 | 2.86951 | ABC transporter ATP-binding protein |
| Q89ZP7 | BT_4329 | -2.97173 | 2.49205 | BFN domain-containing protein |
| Q8A8X2 | BT_1045 | -2.64843 | 2.85322 | Concanavalin A-like lectin/glucanase |
| Q8A7D7 | BT_1587 | -2.61663 | 1.4901 | GCN5-related N-acetyltransferase |
| Q8AAE0 | BT_0525 | -2.32145 | 1.32336 | LruC domain-containing protein |
| Q8A826 | BT_1348 | -2.17801 | 2.67435 | CDP-abequose synthase |
| Q89ZP8 | BT_4328 | -2.167 | 3.63516 | Ribosomal RNA small subunit methyltransferase E |
| Q89ZP9 | BT_4327 | -2.12513 | 5.58384 | DUF4836 family protein |
| Q8A149 | BT_3818 | -1.95031 | 1.38383 | Gliding motility lipoprotein GldH |
| Q89YQ4 | BT_4677 | -1.90211 | 2.01555 | Beta-lactamase-inhibitor-like PepSY-like domain-containing protein |
| Q8A5G3 | BT_2276 | -1.88136 | 1.51383 | AI-2E family transporter |
| Q8A551 | BT_2391 | -1.74464 | 3.18011 | Hybrid Two-component system |
| Q8A6C7 | BT_1957 | -1.70454 | 1.4706 | DUF4465 domain-containing protein |
| Q8A8X3 | BT_1044 | -1.68106 | 3.56162 | Secreted endoglycosidase, GH family 18 |
| Q8A4A5 | BT_2698 | -1.65411 | 1.42781 | Unknown |
| Q8AAL2 | BT_0452 | -1.63773 | 1.32215 | SusC homolog |
| Q89YP0 | BT_4691 | -1.59995 | 1.64528 | Ribosomal RNA large subunit methyltransferase F,rlmF |
| Q8A8X4 | BT_1043 | -1.59626 | 3.77015 | SusD homolog |
| Q8A6F8 | BT_1926 | -1.5911 | 1.87538 | OmpA/MotB domain protein |
| Q8A5H2 | BT_2267 | -1.56716 | 1.5167 | Site-specific integrase |
| Q8A2R4 | BT_3241 | -1.50727 | 1.30956 | SusD homolog |
| Q8A9Y3 | BT_0682 | -1.49005 | 1.74647 | Two-component system response regulator |

| Start Position | End Position | Sequence | Posterior Error Probability |
|---|---|---|---|
| 153 | 164 | EKEEVEPVEETK | 1.62E-05 |
| 187 | 195 | DKLHIPAEK | 0.0033255 |
| 189 | 195 | LHIPAEK | 0.038234 |
| 259 | 279 | QANQVVDMEHHQPISFGLSVR | 3.29E-14 |
| 285 | 302 | GFSVETGLTYTLLSSDAK | 1.01E-36 |
| 314 | 322 | LHYLGIPLK | 6.66E-06 |
| 323 | 330 | ANWNFLDK | 0.0044146 |
| 323 | 331 | ANWNFLDKK | 2.04E-05 |
| 356 | 377 | ETVKPLQFSVSGAVGAQFNATK | 1.58E-09 |
| 378 | 401 | RVGIYVEPGVAYFFDDGSDVQTIR | 4.09E-08 |
| 379 | 401 | VGIYVEPGVAYFFDDGSDVQTIR | 5.89E-08 |
| 402 | 415 | KENPFNFNIQAGIR | 6.49E-12 |
| 403 | 415 | ENPFNFNIQAGIR | 2.13E-09 |

**Table S5: Tryptic peptides of Dma1 identified in the OMV fraction.**
